# Helicon: Helical indexing and 3D reconstruction from one image

**DOI:** 10.1101/2025.04.05.647385

**Authors:** Daoyi Li, Xiaoqi Zhang, Wen Jiang

## Abstract

Helical symmetry is a common structural feature of many biological macromolecules. However, helical indexing and *de novo* 3D reconstruction remain challenging. We have developed a computational method, Helicon, which poses helical reconstruction as a linear regression problem with the projection matrix parameterized by the helical twist, rise, and axial symmetry. A sparse search of the twist and rise parameters would allow helical indexing and 3D reconstruction directly from one 2D class average or a raw cryo-EM image. The Helicon method has been validated with simulation tests and experimental cryo-EM images of helical tubes, non-amyloid filaments, and amyloid fibrils. Imaging stitching and L1 regularization of linear regression were shown to improve the robustness for low-twist amyloids and noisy raw cryo-EM images. Using Helicon, we could successfully index the helical parameters and perform *de novo* reconstruction of a previously unreported, low abundance tau amyloid structure from a publicly available dataset.

**Graphic abstract:** 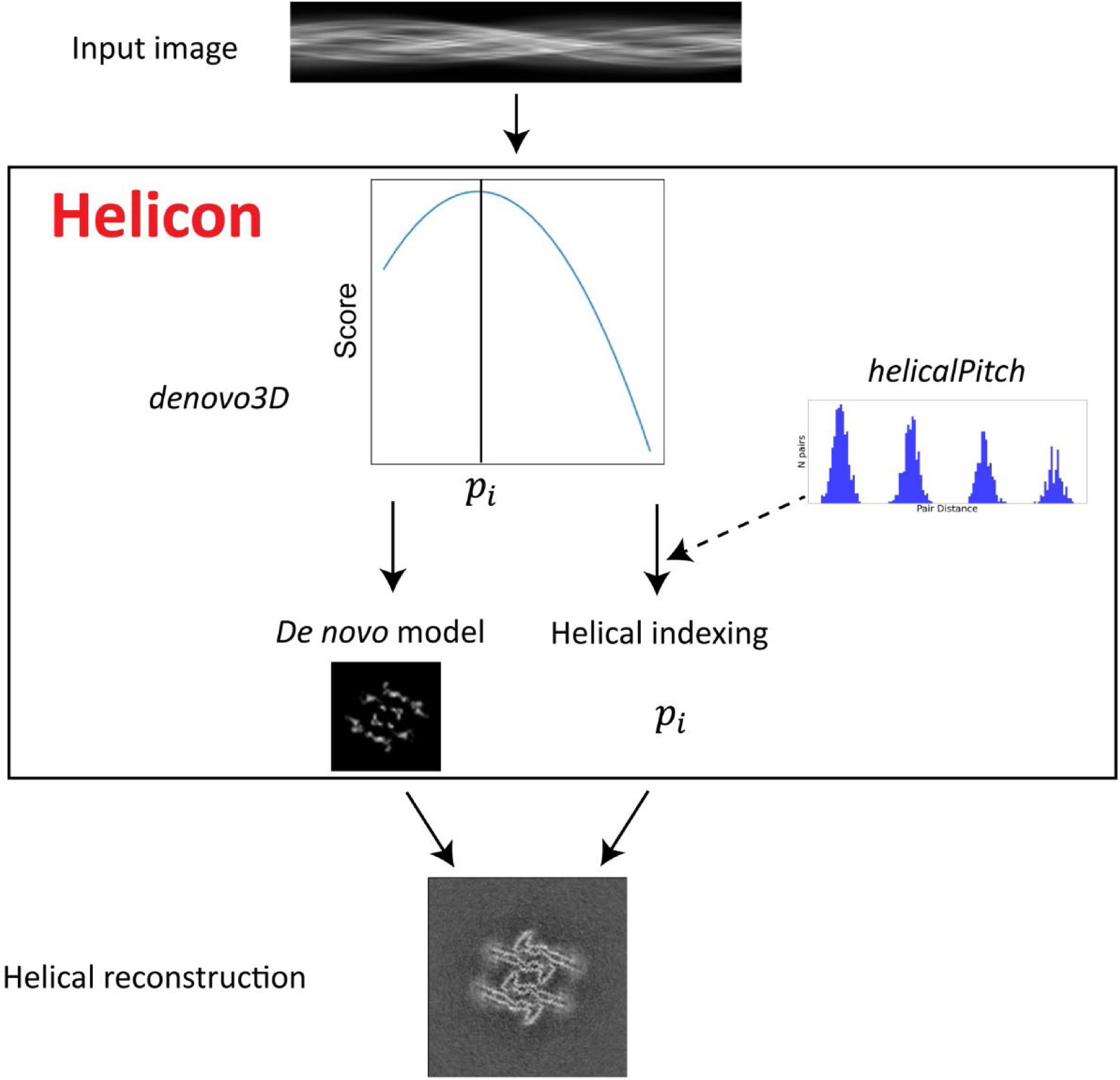

**Highlights:**

- Helicon enables helical indexing and 3D reconstruction from a single 2D image
- Formulates helical reconstruction as a linear regression problem
- Tackles low-twists and raw cryo-EM images with image stitching and regularization
- Validated with diverse experimental data and a previously unreported tau filament

## Introduction

Helical symmetry is a structural feature of many biological macromolecules, including double-stranded DNA^1^, cytoskeletons^2,3^, viruses^4,5^, and pathological amyloid fibrils^6–8^. Determining these helical structures at the atomic level is crucial for understanding their biological functions and can lead to potential drug discoveries^9^. Cryogenic electron microscopy (cryo-EM) is a powerful tool for resolving structures at near-atomic resolutions^10^. Since the first helical structure was solved using EM in 1968^11^, cryo-EM has been widely employed in the structural study of helical structures^12^.

Traditionally, 3D reconstruction of helical polymers is performed in Fourier space using full-length filaments and treating the filaments as 1-D helical crystals. This process involves determining the twist, rise and axial symmetry, by indexing the Fourier-Bessel layer lines and employing Fourier-Bessel synthesis for 3D reconstruction^13^. However, this approach relies on well-ordered helical structures. In contrast, iterative helical real-space reconstruction (IHRSR) treats helical polymers as a series of short, straight segments followed by real-space analysis developed for single-particle reconstructions, with the imposition of helical symmetry^14,15^. Despite this, successful reconstruction of the helical structures from the short segments still requires relatively accurate knowledge of the helical parameters and initial 3D models.

3D reconstruction from 2D projection images requires multiple views of the same molecules, which are typically obtained from different particles for single particle cryo-EM, or from tilt series for electron crystallography and tomography. In contrast, due to the spiraling nature of helical symmetry, unlike typical single particles where each particle image represents a single view, a 2D side projection image of helical structures offers uniformly changing views along the helical axis. This unique property of helical structures allows for 3D reconstruction of the helical structure from a single image if the view angle changes along the axis, i.e. helical twist and rise parameters, are known^16^. For example, the *relion_helix_inimodel2d* program in the RELION software can perform 3D reconstruction from one or multiple 2D class average images with the requirement of a pre-determined crossover value, i.e. known helical indexing^17^.

In this study, we have developed a new method, Helicon, capable of helical indexing and 3D reconstruction of helical structures using a single 2D image, e.g. a 2D class average or a raw cryo-EM image (Fig. 1). Helicon treats the 3D reconstruction problem as a linear regression task *y* = *Ax* in which the voxel values of an asymmetric unit (i.e. *AA*) are estimated from the known 2D image pixel values (i.e. *y*), where *A* is the full projection matrix assembled from the projection matrix of the asymmetric unit and its replicates based on the helical twist, rise, and axial symmetry parameters. A sparse search of the twist and rise parameters would allow helical indexing and 3D reconstruction directly from one 2D image. For amyloids with cross-β stacking of strands with ∼4.75 Å rise along the helical axis, only the twist values in a small range (a few degrees) need to be sparse-searched (e.g. a step size of 0.1°).

**Figure 1:**
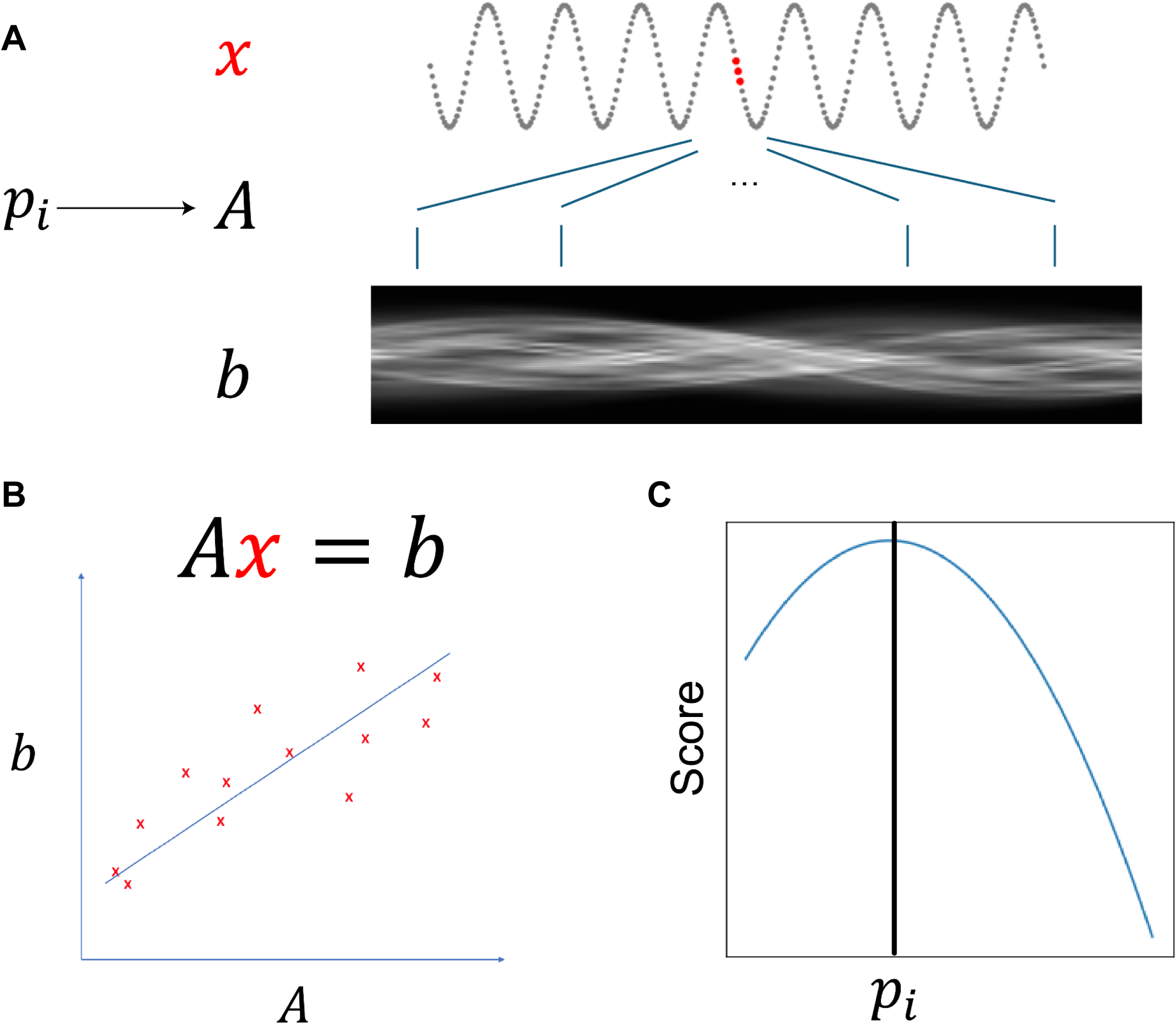
The workflow of the Helicon method. (A) A cryo-EM image *b* of helical structures can be expressed as the projection of a helical 3D density in an asymmetric unit *x* (red) using the projection matrices *A* calculated from the projection matrix of *x* and its replicates transformed with the helical parameters *p*_*i*_. (B) The linear regression model is illustrated using a one-variable example. In the illustration, the unknowns *x* can be solved using the knowns *A* and *b*. (C) Finding the correct helical parameter by raster search of the parameters. Each trial helical parameter will specify a different projection matrix *A* and a corresponding linear regression solution *x*. The parameter *p*_*i*_ and the corresponding solution *x* leading to the highest score will be the final Helicon solution.

Robust, high-quality 3D reconstruction requires full coverage of all views, which is also true for Helicon. While it is generally true for helical structures with relatively short pitches, i.e. a single helical segment containing a half pitch or longer section of the helical lattice, not all helical structures have all the views from a single 2D image, especially for some amyloid fibrils with low twists. For example, an amyloid structure with −0.46° twist and 4.75 Å rise would have a pitch value of 3,800 Å, far larger than the length of typical helical segments^6^. For such low-twist amyloids, a typical image of the helical segment of 200 to 1000 Å in length would only cover a small fraction of the pitch, leading to many missing views. This is similar to the missing wedge problems for tomography^18^. To improve Helicon’s robustness and quality against low-twist helical structures, we have introduced several solutions.

The first approach is regularization, where a penalty term using L1, L2 or both norms is added to constrain the solution values^19^. This regularization ensures that the linear regression model, *x*, will not overfit *y* (i.e. the 2D image), thereby improving robustness. The second approach entails image stitching, whereby images of multiple helical segments are concatenated to generate a single, extended filament image. It could increase the coverage of the helical spiral within a single 2D image and reduce the “missing wedge”, leading to improved robustness of helical parameter estimation and the quality of 3D reconstruction.

The third approach leverages the relative positional information of the helical segments in the parent filaments. This information has been used to classify different filament types^20^ and is now employed to estimate the pitch distance in this work. For this purpose, we developed *helicalPitch*, a Helicon subcommand that analyzes the histogram of pair distances of segments in the same filament being assigned to the same 2D class. The primary peak of this histogram corresponds to the pitch distance of the helical structure represented by this class.

We have validated Helicon using simulated images, 2D class averages, and raw images from experimental cryo-EM datasets, including non-amyloid helical structures and more challenging amyloid fibrils. Helicon thus provides a new approach for helical indexing and *de novo* 3D reconstruction for a wide range of helical structures, including the challenging low-twist amyloids. Additionally, Helicon was able to successfully index and solve the structure of a previously unreported type of tau filament in a publicly available dataset.

## Methods

### 1. Simulation test images

We generated three sets of simulation test images: Gaussian blobs replicated with helical parameters (4.75 Å rise, and 1° twist), projections of non-amyloid helical map EMD-10721 (pili)^21^, and projections of amyloid maps EMD-23871 (*ex vivo* tau)^22^. The EMD-23871 density map was extended to 1000 Å in length based on the helical parameters provided in the EMDB. The maps were downscaled to 5 Å per pixel to mimic the typical box size and pixel size of helical segments used for early rounds of 2D classification^23^.

### 2. Experimental images

We obtained 2D class averages of cryo-EM experimental images of multiple helical structures including non-amyloid and amyloid structures. The images were downloaded from the EMPIAR database: EMPIAR-10019 (VipA/B)^24^, EMPIAR-10640 (α-synuclein)^25^, EMPIAR-10917 (amyloid-β)^26^, and multiple tau datasets with different folds and helical parameters, EMPIAR-10230^27^, EMPIAR-10243^28^, EMPIAR-10940 and EMPIAR-10943^29^. For datasets with deposited 2D class averages (EMPIAR-10940, EMPIAR-10943)^29^, the deposited 2D class averages were used as test images. For other datasets, we reprocessed the data to obtain 2D class averages. Movies were motion-corrected using MotionCor2^30^. The contrast transfer function of all aligned micrographs was estimated using CTFFIND-4.1^31^. For the non-amyloid datasets, filaments were picked using FilamentTracer and further processed with 2D classification in CryoSPARC^32^. For the amyloid datasets which include the particle coordinates in the micrographs, the particles were extracted from the micrographs using the deposited coordinates. The 2D classification was then performed in RELION with the first peak of the CTF ignored to preserve more low-resolution information^15^.

For Helicon 3D reconstruction directly from raw images, we used the type 3b amyloid-β filaments in the *ex vivo* amyloid dataset of a Down syndrome patient^6^ and the 2^nd^ type of tau filaments in EMPIAR-10940. We pre-processed the movies in EMPIAR-10940 using the same procedure as described above. The raw filament images were cropped from the micrographs using the *e2helixboxer.py* and low pass filtered to 20 Å with *e2proc2d.py* in the EMAN2 software^33^.

### 3. Linear regression model

We use *ρ*(*r*, *φ*, *z*) to represent a helical structure in cylindric coordinate with *r* for the radius from the helical axis, *ϕ* for the rotation angle around the helical axis, and *z* for the position along the axis. For a helical structure with a twist of Δ*ϕ* and an axial rise Δ*z*, we should have:

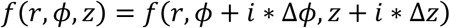

where *i* is an integer. This indicates that a helical structure is invariant to the shift and rotation corresponding to the helical parameters. If we take the volume of one asymmetric unit of the helical structure as *x*, the helical structure *V* is represented as:

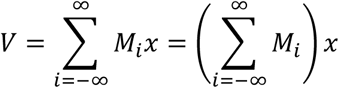

where *M*_*i*_ is the rotational and translational transformation corresponding to the *i*-th integer multiple of the helical twist Δϕ and rise Δz, with *i* ∈ [−∞, ∞]. *M*_*i*_ can be represented as a 4×4 transformation matrix:

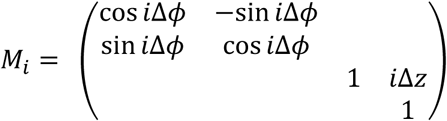

In this notation, the helical axis is along the Z-axis. As the elongated helical structures lie in the x-y plane of the grid during data collection, the helical structure *V* will be tilted 90° (*M*_90_) before being projected along the *z* direction to obtain the final projected image *I*:

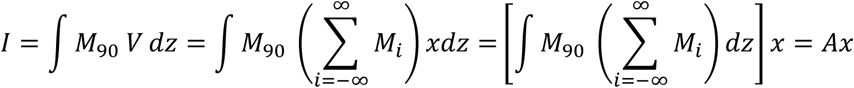

The projection *I* is expected to match the final targeted image *b*, i.e. we should have

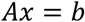

if the *A* and *x* are both correct (Fig. 1). This equation is a linear regression problem that can be solved by minimizing the following errors:

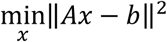

Since *A* is parameterized by the helical parameters twist and rise, we would need to search for a range of twist and rise values, build the *A* matrix for each set of helical twist/rise, and solve the linear regression problem to obtain the solution *x* (i.e. the voxel values of the helical asymmetric unit). The optimal twist/rise is the one that produces a projection most closely resembling the target image. Thus, we can calculate the correlation between the projected image (*Ax*) and the target image (*b*) as the following to find the optimal twist/rise that maximizes the similarity score:

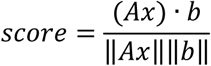

### 4. Regularization

When the length of the helical segment exceeds 0.5x pitch, i.e. the tilt angle covers at least 180° or −90° to 90° tilt range, an ordinary linear regression solution is effective. However, for low-twist helices where the helical segment length is less than 0.5x pitch, the problem becomes analogous to tomography with missing wedge or single particles with preferred orientations, resulting in insufficient angular information to represent the projection accurately. Consequently, ordinary linear regression solutions are susceptible to overfitting and poor solutions.

To mitigate this problem, we employ the ElasticNet regression model to introduce regularization into the loss function ^19^. The ElasticNet combines both L1 and L2 norms as the regularization penalties, which helps prevent overfitting by balancing between the penalties of these two norms. The objective is to minimize the following equation:

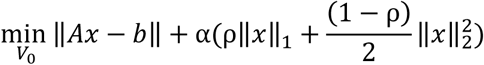

where *α* is the weight of the regularization of the ElasticNet and *ρρ* represents the relative weight between the L1 regularization and L2 regularization.

### 5. Web app user interface

The linear regression formulation assumes that the 2D images of helical structures are horizontal (i.e. along the X-axis) and centered in the Y-direction while the 2D class averages generated by popular programs such as RELION and CryoSPARC are often not ideally aligned. To help mitigate this issue, we have developed the above linear regression method as *helicon denovo3D*, a Web app with a graphical user interface (GUI) to facilitate the centering and horizontal alignment of helical 2D images for Helicon. This GUI interface allows users to transpose the CryoSPARC 2D class averages oriented in Y-direction to align with the X-axis, and manually adjust in-plane transformation parameters, ensuring that the helical axis aligns with a centered, horizontal reference line. The user-friendly GUI provides an effective solution for handling 2D class averages that exhibit misalignment (Fig. S1).

However, for 2D class averages with out-of-plane tilt, simple in-plane transformations are insufficient for proper correction. Despite this limitation, Helicon can achieve meaningful 3D reconstructions and accurate helical parameter estimations for images with tilts of up to 10 degrees (Fig. S2). To further understand the severity of out-of-plane tilts, we performed simulation experiments with grids of different out-of-plane tilt angles and filaments having random in-plane directions. The histograms of tilt angles of the helical segments peak at 90 degrees (Fig. S3). This suggests that most filaments in cryo-EM datasets would only exhibit minor out-of-plane tilts, within the working tilt range of Helicon (Fig. S2).

To overcome the low twist challenges, we have included image stitching in the Web GUI. The user can choose multiple images and interactively adjust their in-plane rotations and positions to superimpose them, ensuring that overlapping regions between different images are properly aligned (Fig. S4). Subsequently, we employ the ITKMontage package^34^ to merge the images into a single composite image.

### 6. *HelicalPitch* Web app

To address the challenges with low twist helical structures, where a single 2D class average cannot provide sufficient information for robust *de novo* identification of helical twist by Helicon, we have developed the *helicon helicalPitch* Web app to complement the *helicon denovo3D* Web app. In this method, we leverage the metadata of extracted helical segments (e.g. parent filament ID, location in the micrograph) and the 2D classification results (e.g. class ID). We compute the pair distances among all segments in the same parent filament being assigned into the same 2D class and then plot the histogram of the pair distances. Peaks with uniform spacing are expected in this plot. The first major peak in the histogram plot corresponds to the distance at which the helix has spiraled along the helical axis to repeat within the filaments. This distance (*Pitch*^∗^) is related to the pitch and the axial symmetry (n_sym_) of the helical structure, and can be expressed by the following formula:

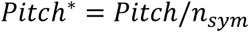

### 7. Implementation

The Helicon method was implemented in Python utilizing the linear regression models in the scikit-learn library. The GUI of *helicon helicalPitch* and *helicon denovo3D* were implemented as Web apps using the py-shiny library. Helicon is openly developed, with its source code and updates available on GitHub (https://github.com/jianglab/helicon). Helicon also includes several useful tools as sub-commands, for example, *images2star* and *cryosparc*, which help users read/convert/transform RELION star files and CryoSPARC cs files, respectively. Detailed descriptions of these additional functionalities are beyond the scope of this publication and the readers are encouraged to download and explore.

1. 8. Helical reconstruction of the second type of tau filament unreported from EMPIAR-10940

The 2^nd^ type of filament in EMPIAR-10940 was first visually identified from the 2D class averages. The helical twist was determined by the consensus result of Helicon *denovo3D* from the raw images and 2D class averages and the result of *helicalPitch* using the 2D classification metadata. The initial 3D map was reconstructed from the raw images using *denovo3D* with the consensus twist parameter. Several rounds of 3D classification were carried out to obtain the best homogeneous subset. A full-filament recovery strategy was applied after every round of selection^20^. The final selected segments were used for 3D auto-refinement with blush regularization^35^ yielding a 3D map showing well-resolved backbone and side-chain densities.

## Results

### 1. Validation using simulation images

We first validated the *Helicon denovo3D* algorithm on simulation images containing projections of Gaussian blobs. The *denovo3D* algorithm successfully determined the helical parameters, as shown by the helical parameters with the highest score corresponding to the correct parameters (Fig. 2A). Additionally, the reconstruction was visually identical to the ground truth density (Fig. 2D, S5A,E). While *helicon denovo3D* allows the users to specify the axial symmetry, we only used the default C1 symmetry to avoid imposing prior knowledge. The axial symmetry recovered by Helicon is thus an unbiased result that can be used as a validation criterion for the Helicon reconstruction. As shown in Fig. 2D, the 2-fold symmetry in the test structure was recovered by Helicon. We then applied *helicon denovo3D* to the projection image of a non-amyloid helical structure, bacterial pili, generated from the extended 3D density map of EMD-10721 ^21^. Helicon accurately determined the helical parameters of the pili and successfully reconstructed the structure (Fig. 2B,E). The FSC of the reconstructed density and the original density remains higher than 0.5 at the Nyquist (10 Å) of the binned volume (Fig. S5B,F). Therefore, we also tested Helicon using an un-binned projection image with a pixel size of 0.87 Å. The FSC curve reports 4.9 Å at the 0.5 threshold (Fig. S5C,G).

**Figure 2.**
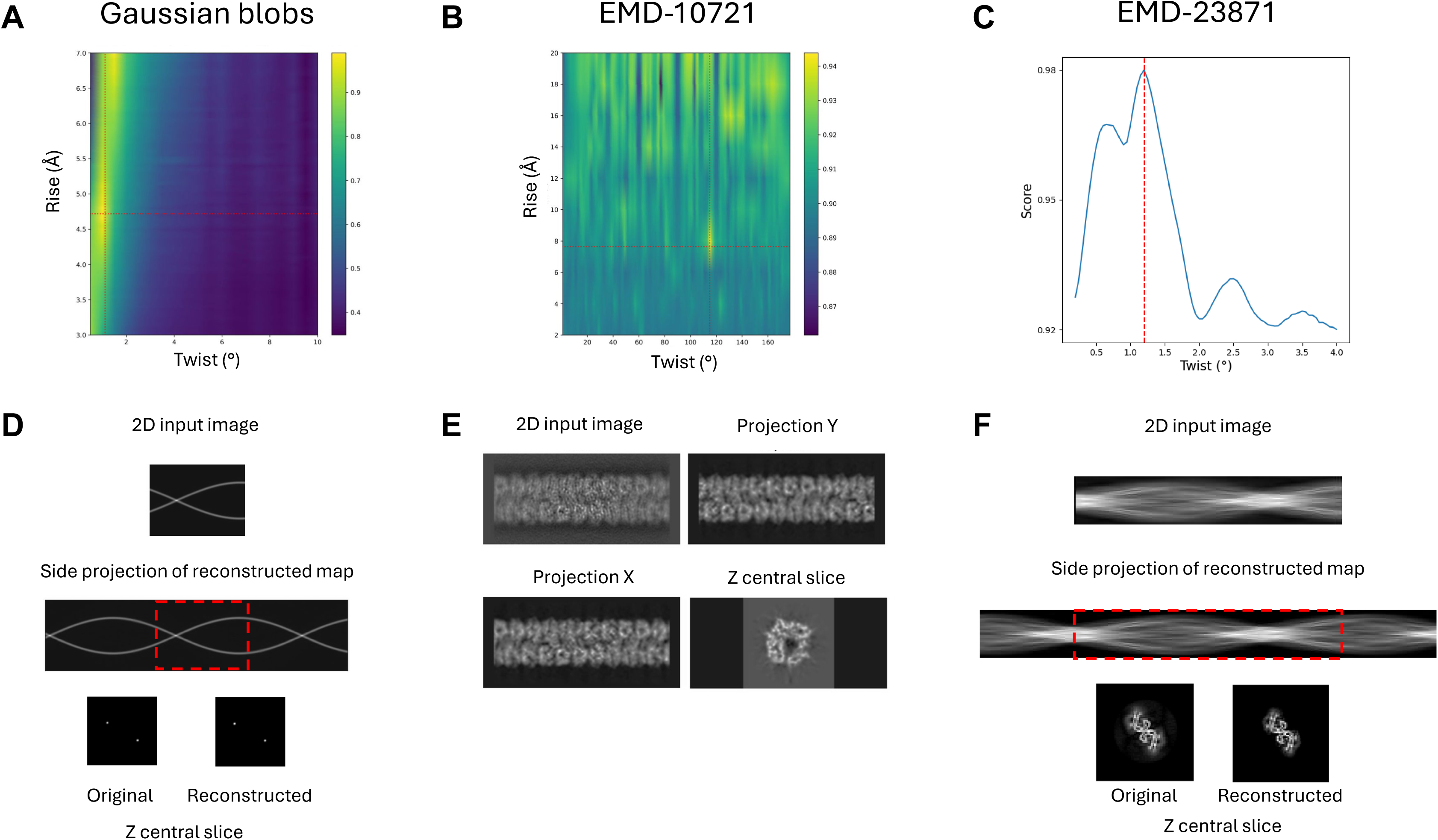
Validation of Helicon using simulated images. (A-C) Scores of Helicon validation tests using 2D projections of simulated Gaussian blobs (A), bacterial pili (EMD-10721) (B), and PHF tau filaments (EMD-23871) (C). Red dashed lines indicate the ground truth values of the helical parameters. (D-F) The input test image, projections, and central Z slice of Helicon 3D reconstructions using the helical parameters that produced the best score as shown in A-C, respectively. Red boxes denote the reconstructed regions corresponding to the original input images.

For non-amyloid helical structures, the extracted segment images are typically longer than one pitch, providing full coverage of views around the helical axis to solve the linear regression problem without overfitting. As amyloid structures exhibit much smaller twists (e.g. up to a few degrees), it is common that the 2D segment images only cover a helical section much smaller than one pitch, which will potentially lead to overfitting of the linear regression model. Therefore, in the simulation test of amyloid structures, we extended the EMDB density of the paired helical filament (PHF) tau to 1000 Å based on the provided helical parameters, approximating half of the pitch of PHF tau structure, and generated the projections. *Helicon denovo3D* could also accurately determine the helical twist of the amyloid structure (Fig. 2C), and the reconstruction quality was excellent as shown by the FSC with the ground truth map staying above 0.5 across the entire spatial frequency range (Fig. 2F, S4D,H).

### 2. Testing using 2D class average images of non-amyloid helical structures

After validating the *denovo3D* algorithm with satisfactory reconstruction quality using simulated projection images, we tested cryo-EM experimental data using 2D class averages of non-amyloid helical structures. Based on the morphology of the helical structures, they were grouped into two categories. The first group included tubular structures with larger diameters and hollow interiors, such as VipA/B and TMV structures. The second group comprised filamentous structures, such as bacterial pili, which are typically thinner. For the first group, using the 2D class average of the VipA/B images from EMPIAR-10019^24^, *Helicon denovo3D* accurately determined the helical twist and rise parameters and successfully reconstructed the 3D density (Fig. 3A,C, S6A). For the second group, using the 2D class average of bacterial pili images, *Helicon denovo3D* also accurately determined the helical twist and rise parameters and reconstructed the helical structure (Fig. 3B,D, S6B).

**Figure 3:**
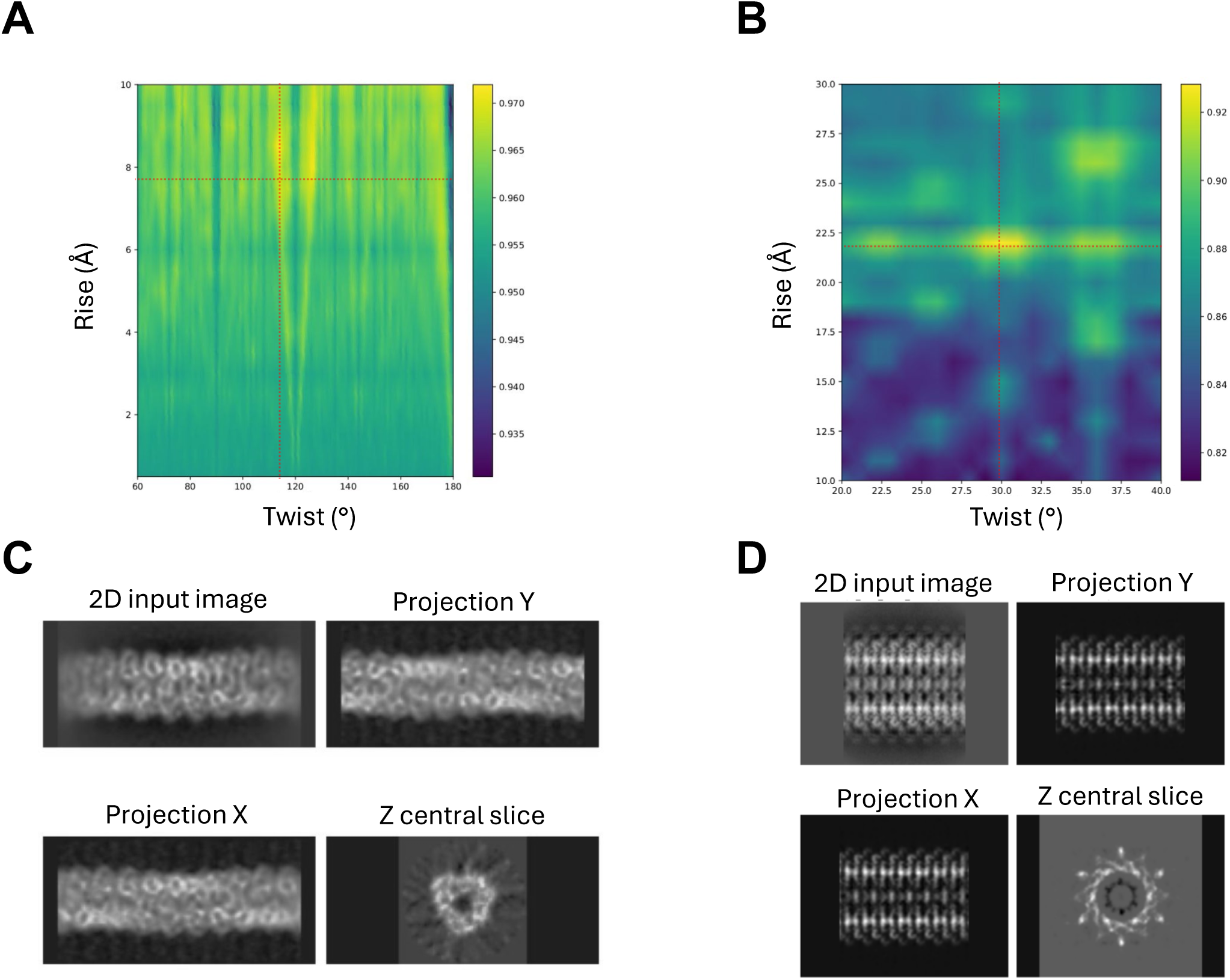
Testing Helicon using 2D class averages of non-amyloid helical structures. (A, B) Scores of Helicon tests using 2D class averages of bacterial Pili (A) and VipAB (EMPIAR-10019) (B). (C, D) The input 2D class average images, X, Y projections, and central Z slice of Helicon 3D reconstructions using the helical parameters yielding the best score as shown in A and B, respectively. Red dashed lines mark the ground truth values of the helical parameters.

### 3. Testing using 2D class average images of amyloid helical structures

*Helicon denovo3D*’s performance on non-amyloid helical structure images was promising, prompting us to test it on amyloid structures. We first tested it on images of recombinant tau amyloids. Compared to datasets of *ex vivo* samples extracted from brain tissues, recombinant tau protein datasets have fewer contaminants and no fuzzy coat of disordered peptide regions, significantly enhancing the quality of 2D classification^20^. The 2D averages were obtained directly from the EMPIAR repository^29^. Helicon’s determined helical parameters were consistent with the deposited values in the EMDB (Fig. 4A), and the reconstructed map matched the EMDB density (Fig. 4D). To verify the robustness of this method, we also tested it on another cryo-EM dataset of recombinant amyloid α-synuclein fibrils^25^. Helicon determined the helical twist with a clear peak value consistent with the deposited twist value in the EMDB (Fig. 4B), and the reconstructed map matched the EMDB density (Fig. 4E).

**Figure 4:**
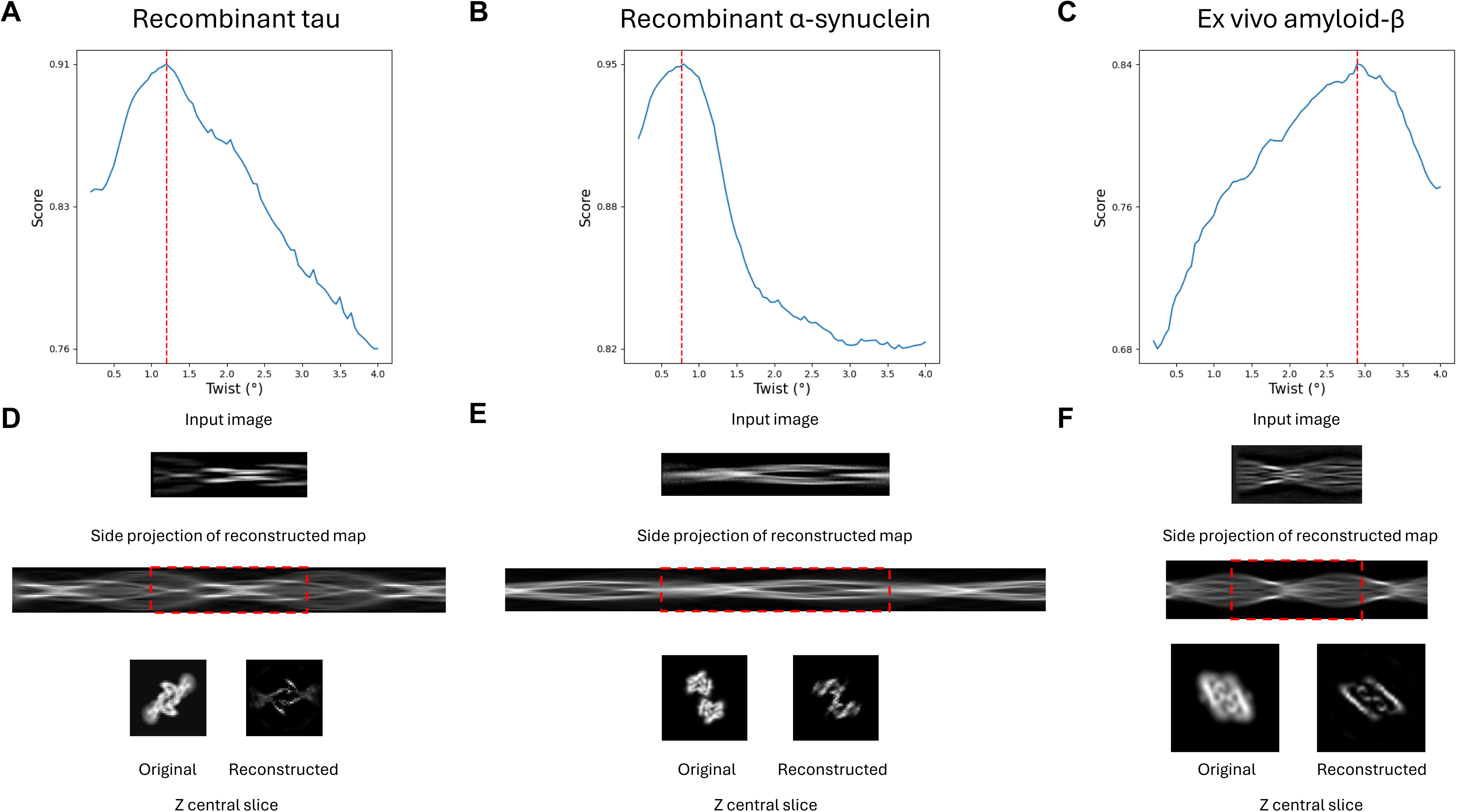
Testing Helicon using 2D class averages of amyloid helical structures. (A-C) Scores of Helicon tests using cryo-EM 2D class averages of recombinant tau in EMPIAR-10940 (A), α-synuclein filaments in EMPIAR-10640 (B), and amyloid-β fibrils in EMPIAR-10917 (C). Red dashed lines indicate the ground truth values of the helical twist (D-F) The input 2D class average images, side projection, and central Z slice of Helicon 3D reconstructions using helical parameters that provide the best score as shown in A-C, respectively. Red boxes denote the reconstructed regions corresponding to the original input images.

We then tested *helicon denovo3D* on cryo-EM images of an *ex vivo* amyloid-β sample extracted from human brains, which contained more contaminants^26^. After performing 2D classification of the extracted helical segments, we selected the 2D class with the most visual features for the Helicon test. The helical parameters were correctly determined (Fig. 4C) for this amyloid-β structure and the morphological features of the reconstructed amyloid-β structure matched the corresponding EMDB density (Fig. 4F).

### 4. Tackling the low twist amyloid challenges with regularization of the regression model and image stitching

The results above demonstrate that *helicon denovo3D* performs well on 2D images covering more than 0.5x pitch (i.e. one crossover when the helical structure has an elongated shape in XY plane, resulting in visually obvious changes of the diameter along the fibril), where projections of the helical asymmetric unit are available from all views around the helical axis^6^. When working with low-twist amyloids, the helical segment length is often shorter than 0.5x pitch. This is equivalent to the preferred orientation problem in single particle cryo-EM or missing wedge problem in electron tomography or crystallography, causing the linear regression model to overfit the limited data. Consequently, the model with the best score and the corresponding helical parameters may be incorrect (Fig. 5A,D).

**Figure 5:**
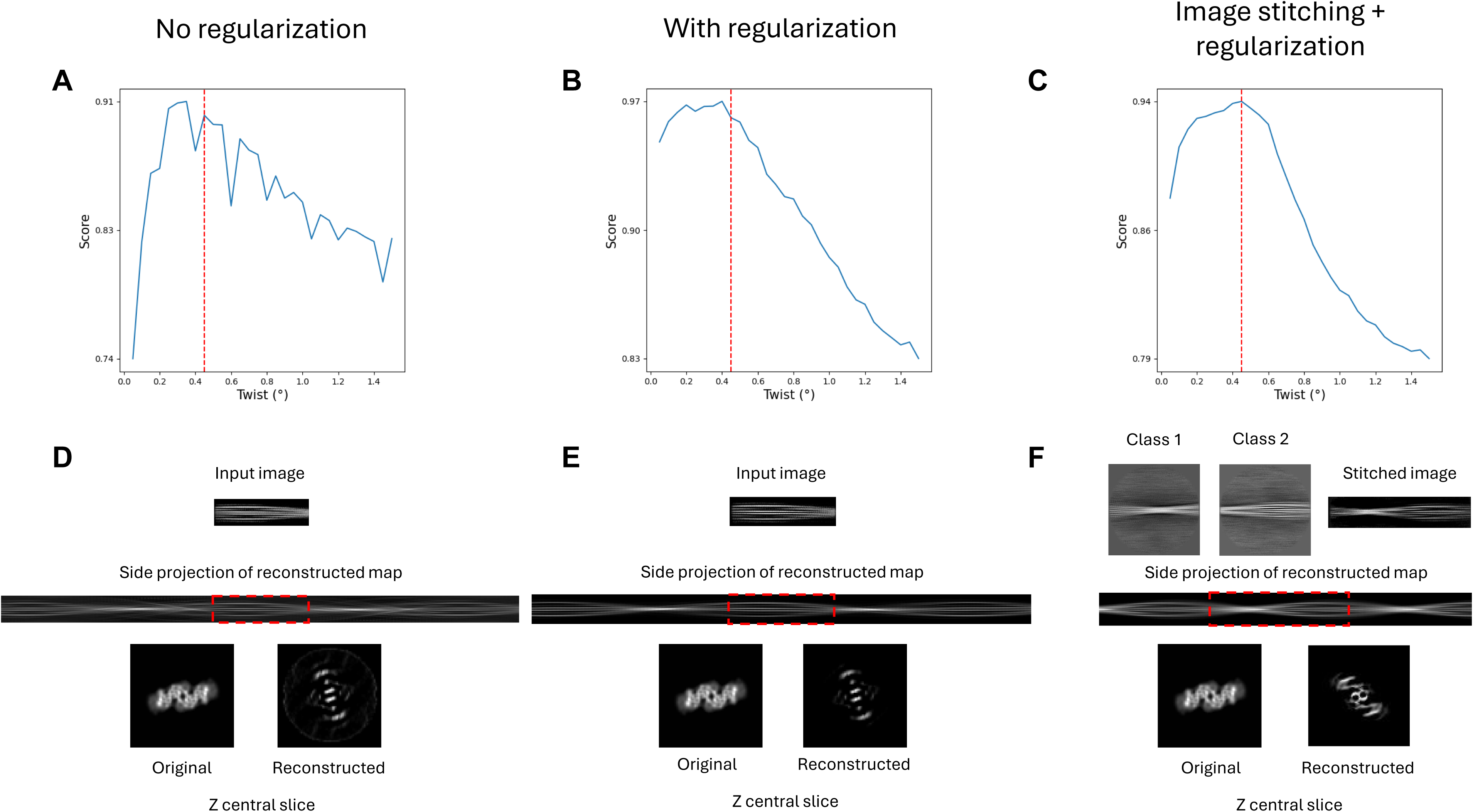
Improving Helicon solutions for low-twist amyloid structures using regularization and image stitching. Class averages of type 3b amyloid-β fibrils in brain samples of Down Syndrome patients were shown in this example. (A-C) Scores of Helicon solutions without regularization (A), with regularization (B), and with both regularization and image stitching (C). (D-F) The input class average image, side projection, and central Z slice of Helicon 3D reconstruction using the helical parameters that achieved the best score in each experiment shown in A-C, respectively. Red dashed lines indicate the ground truth value of the helical twist, and red dashed boxes denote the reconstructed regions corresponding to the original input images.

To address the low twist challenges, we introduced an ElasticNet regularization term to the linear regression model to mitigate overfitting. Figure 5B shows an example using a 2D class average (∼1,600 Å image box size) of cryo-EM data of the type 3b Aβ that has a −0.46° twist and 3,725 Å pitch ^6^. Without regularization, incorrect helical twist yielded the best score, indicating overfitting of the linear regression model (Fig. 5A). The corresponding reconstructed helical density was noticeably incorrect (Fig. 5D). By introducing ElasticNet regularization, the helical twist with the best score approached the ground truth using an appropriate regularization strength (Fig. 5B, S7). However, the reconstruction quality remained suboptimal due to insufficient view coverage (Fig. 5E).

To further improve the reconstruction quality, we explored the concatenation of multiple 2D images to increase the effective length and view coverage of the stitched segment. We started with the recombinant tau images, which do not have the low twist problem. Our tests showed that stitching images into a longer single image sharpened the peak of the best *denovo3D* score and confirmed that the reconstructed helical structure stayed consistent with the ground truth (Fig. S8A,B). The reconstruction also matched the ground truth more closely than the reconstruction using the shorter 2D average image (Fig. S8C,D). Applying stitching and regularization to the challenging, low-twist type 3b Aβ image, the best helical twist found by Helicon became the same as the ground truth value. More excitingly, the reconstruction quality improved dramatically, capturing most of the structural features in the ground truth density (Fig. 5F).

### 5. Testing using raw cryo-EM images

*Helicon denovo3D* has so far demonstrated satisfying performance using clean 2D class average images. We then tested it with simulated noisy images and raw cryo-EM images to gauge the robustness of Helicon against noise and the feasibility of 3D reconstruction using raw images. Direct reconstruction using raw images provides multiple benefits including avoiding potential errors and artifacts in 2D classification^20^ and using long, raw filaments to better address the challenges with low-twist amyloids.

The tests with simulated noisy images (Fig. S9) allowed us to further characterize the importance of regularization, especially the L1 regularization. We observed that L1 regularization yielded more accurate estimations of the helical twist value at high noise levels, likely due to its tendency to promote sparsity in the solutions^36^. This sparsity is particularly advantageous for noisy data, as it enhances the ability to extract meaningful structural features from degraded signals, thereby improving reconstruction accuracy^37^. Conversely, L2 regularization empirically resulted in superior reconstruction quality using raw cryo-EM images, where more details are preserved compared to the results using L1 regularization (Fig. S9). Based on these findings, we adopted a hybrid strategy: employing a linear regression model with L1 regularization for helical parameter estimation, followed by L2 regularization for reconstruction. The Helicon Web app allows the users to easily choose the relative weights of L1 and L2 norms of ElasticNet regularization, L1-only (Lasso), or L2-only regularization.

We then tested *Helicon denovo3D* on raw cryo-EM images of two recombinant tau datasets and one *ex vivo* Aβ amyloid dataset, where the last one was the low-twist type 3b Aβ amyloid. The raw filaments were first 20Å low pass filtered to reduce noise and enhance the low-resolution morphology features. The helical parameters estimated by Helicon for the raw images of the two recombinant tau datasets were consistent with the values deposited in the EMDB (Fig. 6A,B). The reconstructed density maps demonstrated a high degree of agreement with the corresponding EMDB reference densities, including the fold and the axial symmetry (Fig. 6D,E). More excitingly, Helicon could reliably identify the correct twist value of the low-twist type 3b Aβ filament in an *ex vivo* dataset (Fig. 6C). Furthermore, the reconstructed density closely resembles the deposited density in the EMDB (Fig. 6F).

**Figure 6:**
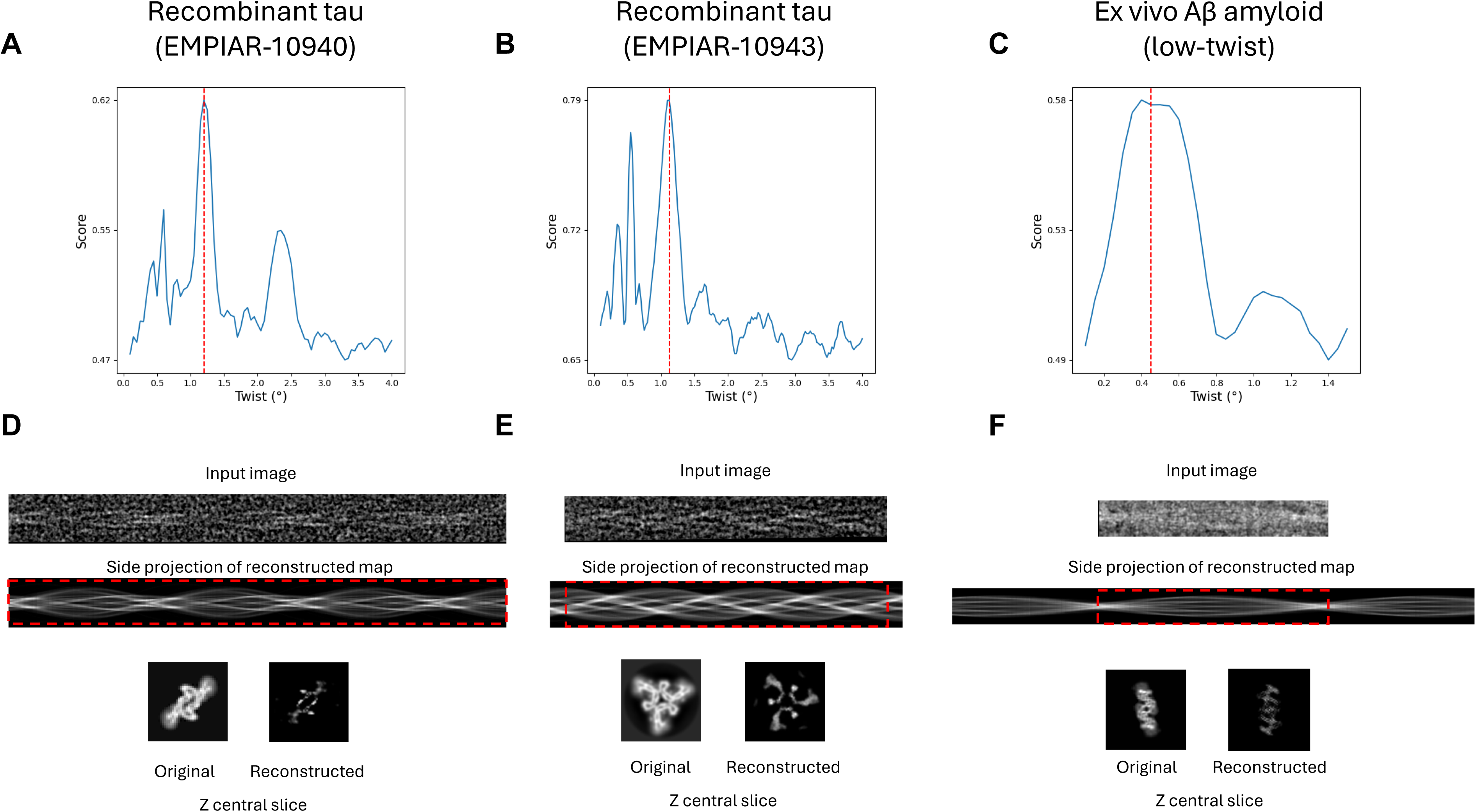
Testing Helicon using raw cryo-EM images. Scores of Helicon tests using the raw cryo-EM images of recombinant tau in EMPIAR-10940 (A), EMPIAR-10943 (B), and *ex vivo* Aβ in Down syndrome sample (C). Red dashed lines indicate the ground truth values of the helical twist (D-F) The input raw cryo-EM images (low-pass filtered to 20 Å), side projection, and central Z slice of Helicon 3D reconstructions using helical parameters that provide the best score as shown in A-C, respectively. Red boxes denote the reconstructed regions corresponding to the original input images.

### 6. Estimating helical pitch using 2D classification metadata

The Helicon linear regression model has demonstrated sufficient performance in estimating the helical parameters and reconstructing 3D models from 2D images for a wide range of helical structures. However, its performance, particularly regarding reconstruction quality, diminishes when applied to low-twist amyloid structures. It benefits from another perspective to obtain more evidence that cross-validates the *Helicon denovo3D* solution. An orthogonal source of information that remains unutilized is the relative positional information of the helical segments in current image analysis workflows for helical structures. Here, we correlated the relative positions of helical segments with the class assignments from 2D classification by calculating the distances between all pairs of helical segments within the same parent filaments assigned to the same 2D class. In the histogram plot of the pair distances for one or more 2D classes, we hypothesize that equally spaced peaks should be present, and the position of the first non-zero primary peak should be equal to the helical pitch divided by the axial symmetry (Fig. 7A). The axial symmetry can often be visually inferred from the 2D class averages. For instance, C1 symmetry does not exhibit clear symmetrical features in the 2D class average (Fig. 7B). In contrast, C2 and 2_1_ symmetries present vertical mirror symmetry in the image (Fig. 7C,D). C3 symmetry exhibits staggered vertical mirror symmetry (i.e., vertically flipping the image and then shifting along the axis to match the original image) (Fig. 7E). For higher-order symmetries, visually determining the axial symmetry from the 2D class average becomes more challenging. Once the axial symmetry is identified, the pitch value can be calculated, and with the helical rise at ∼4.75 Å for amyloids, the corresponding twist value can be derived.

**Figure 7:**
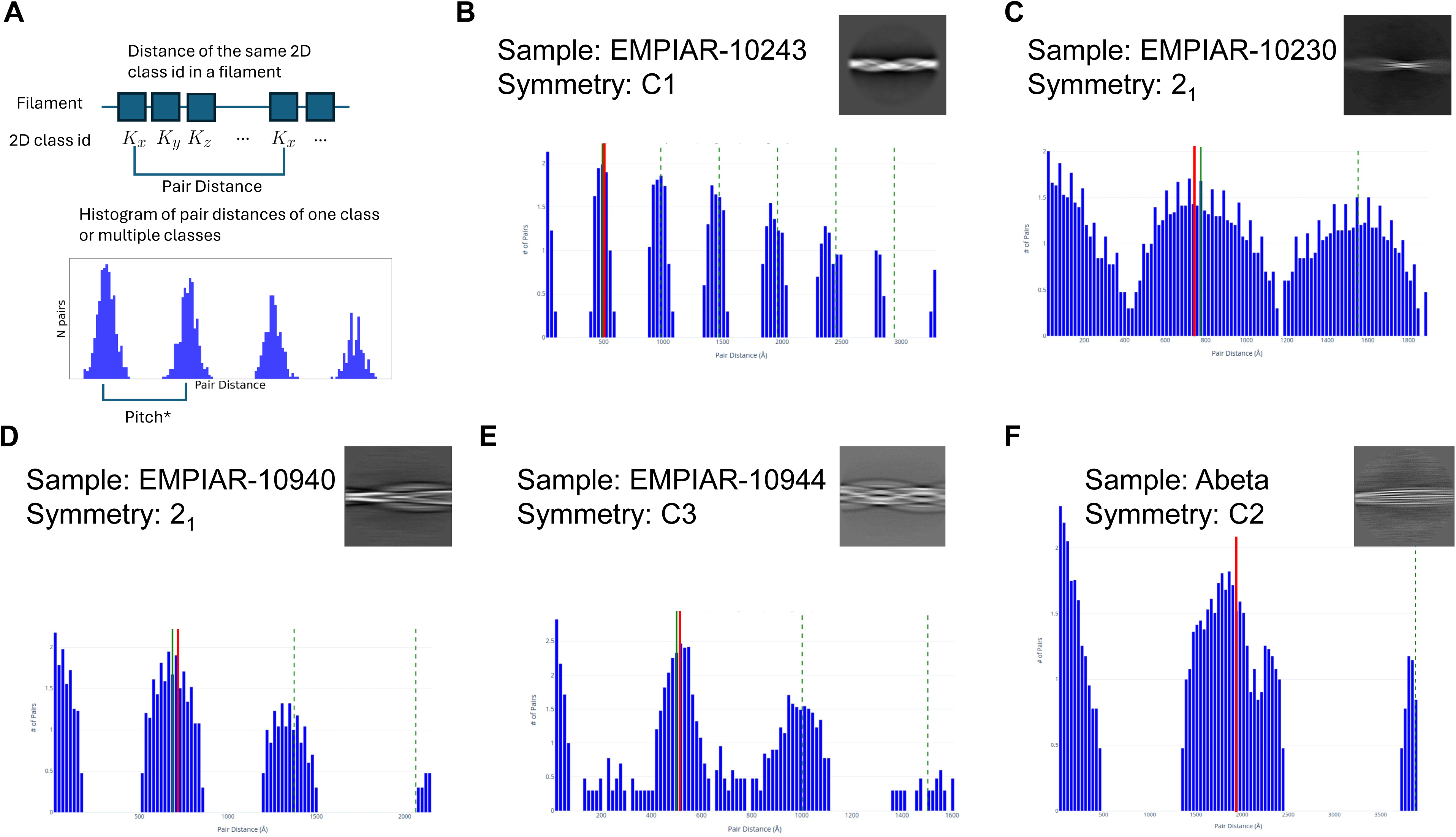
Estimating helical pitch using the relative distances of pairs of segments assigned to the same class by 2D classification. (A) An illustration of the *helicalPitch* approach. (B-F) Histogram plots of the pair-distances of helical segments in the same parent filaments assigned to the same class by 2D classification of cryo-EM images of *in vitro* tau from EMPIAR-10243 dataset with C1 symmetry (B), *ex vivo* PHF tau with 2_1_ symmetry in EMPIAR 10230 (C), *in vitro* tau with 2_1_ symmetry from EMPIAR-10940 (D), *in vitro* tau with C3 symmetry from EMPIAR-10943 (E), *ex vivo* type 3b amyloid β with C2 symmetry from the Down syndrome patient brain samples (F). In panel F, the plot includes the pair distances for multiple classes, and only filaments longer than 3000 Å and multiple 2D classes are selected. Red line: the ground truth pitch value accounted for the axial symmetry. Solid green line: the estimated pitch* value. Dashed green line: integer multipliers of the estimated pitch* value.

To validate this idea, we implemented it in the *helicon helicalPitch* Web app and tested it with multiple experimental datasets. We initially applied it to cryo-EM amyloid 2D class averages that do not suffer from low-twist problems. Our analysis confirmed that this method can reliably determine the twist value from the clear peaks with regular spacing in the pair-distance histograms for both *in vitro* and *ex vivo* amyloid structures across different axial symmetries (Fig. 7B-E). We then applied this method to the low twist 2D class shown in Figure 5. However, the histogram plot was broad and did not provide a clear peak to allow reliable estimation of the pitch distance when including all filaments in the plot (Fig. S10A). Due to the low-twist nature of this helical structure, the pitch is expected to be relatively long, and short filaments, if shorter than a pitch*, are unlikely to provide meaningful pair information. Pair distances from those short filaments are thus most likely to contribute false pair information and potentially introduce noise. Therefore, we restricted our analysis to long filaments within the dataset (Fig. S10B). However, the number of long filaments was limited, so it was necessary to combine the pair distances of multiple 2D classes on long filaments in the histogram plot. With this optimization, the histogram was noticeably improved, and the pitch value could be reliably estimated (Fig. 7F). Automated filament length selection is thus enabled as default in the *helicalPitch* Web app.

### 7. *De novo* indexing and 3D reconstruction of a second type of tau filament structure using Helicon

During the reprocessing of the EMPIAR-10940 recombinant tau dataset, we unexpectedly noticed 2D classes with a larger diameter and a morphology distinct from that of the previously reported type^23,29^ (Fig. 8A). Mapping the classes to raw micrographs confirmed the 2^nd^ type of filaments that has not been reported in this dataset (Fig. 8B). We thus used this filament as a blind test for Helicon to perform *de novo* indexing and 3D reconstruction. Using raw filament and 2D class images, the twist search using *helicon denovo3D* consistently resulted in a clear peak at 1.0° and corresponding 3D models (Fig. 8C,D) resembling a doublet of the previously reported structure (EMD-14046) of the more abundant type of filaments in this dataset. While these consistent results appear convincing, the small difference in the twist angle (1.0°) from the 1.2° twist of EMD-14046 initially raised some concerns about the accuracy of *helicon denovo3D*. However, the concern is mitigated by the same 1.0° twist found by *helicon helicalPitch* based on orthogonal information (Fig. 8E). The *helicon denovo3D* reconstruction from the raw filament image was subsequently used as the initial model for further RELION refinement, which confirmed the −1.0° twist and resulted in a final map at 4.24 Å resolution displaying well-resolved densities for the main chain and visible side chain densities (Fig. 8F, S11A). We subsequently found that this amyloid fold with four protofilaments, although not previously reported for this dataset, was reported in a condition using a different tau protein fragment^29^. It was interesting to observe that the 3D reconstruction of this 2^nd^ type of filament was accomplished using only 2,172 helical segments extracted from 44 filaments. When only the two longest filaments (343 segments) were used, the structure could be solved to 4.93 Å resolution with only slightly reduced structural details (Fig. 8F, S11B). Since both reconstructions used very small datasets due to the low abundance of this filament type, some fluctuations in the FSC curve were expected. (Fig. S11).

**Figure 8:**
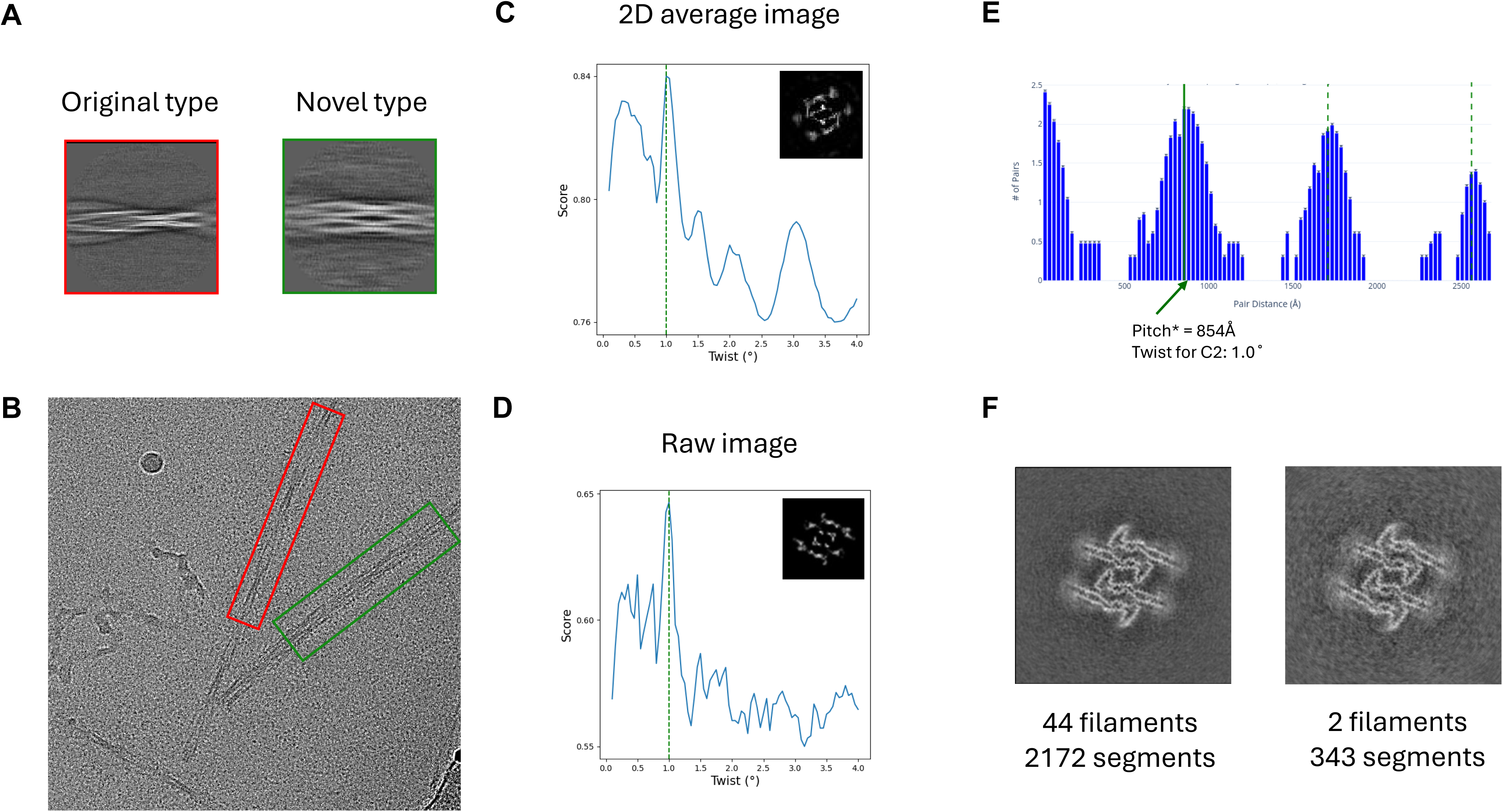
*De novo* indexing and reconstruction of a 2^nd^ type of tau filament using Helicon. (A) Representative 2D classes of recombinant tau filaments in EMPIAR-10940. Left with red box: the previously reported type (EMD-14046); Right with green box: a 2^nd^ type of filament identified in this work, unreported in this dataset. (B) A micrograph in which both types of tau filaments are marked, red box for the previously known type and green box for the 2^nd^ type of filament, respectively. (C, D) Helicon twist search scores using a 2D class average (C) and the raw filament image shown in (B). Green dashed lines indicate the estimated twist value represented by the best score. The central Z-section of Helicon reconstructions are shown as insets. (E) The *helicalPitch* result of the 2D class of the 2^nd^ type of tau filaments. The green line indicates the pitch* value. (F) The final refinement results for the 2^nd^ type of tau filament using either all 44 filaments or only the 2 longest filaments. The central Z-section of the final density maps are shown. The Helicon 3D model from the raw filament image (D) was used as the initial model for both refinements.

## Discussion

In this study, we explored the feasibility of using a linear regression model to obtain 3D reconstructions of helical structures using a single cryo-EM 2D image. The inherent advantage of helical symmetry, which provides uniformly varying views along the helical axis, underpins the effectiveness of this approach. Helicon was extensively validated on simulated images, and demonstrated success with 2D class averages of multiple experimental cryo-EM images, including low-twist amyloids with pitch lengths far exceeding the length of the helical segments. Helicon was also validated on raw filament images directly extracted from micrographs, signifying its robustness and applicability to experimental data.

One significant challenge is the reconstruction of low-twist amyloid structures such as type 3 Aβ^6^. These structures present inherent difficulties due to the limited coverage of view angles by the 2D class averages of helical segments. While our study showed that appropriate levels of regularization can improve the accuracy of helical parameter estimation, this approach using regularization does not fully resolve the issue. We also tackled the low-twist problem from another orthogonal perspective, which is to utilize the relative positional information of helical segments extracted from the same parent filaments. Results from both approaches can be used to cross-validate each other and improve the reliability of the helical parameter estimations.

The lack of sufficient angular information in low-twist helical structures is analogous to the preferred orientation problem in single-particle analysis^38^. Consequently, although helical parameters can be estimated with improved accuracy using regularization or the *Helicon helicalPitch* program, the reconstructed structures might still lack meaningful details, even when the correct helical parameters are provided. To mitigate this limitation, we have incorporated image stitching, where multiple 2D images with overlapping angular coverage are concatenated to form an extended helical image. This approach effectively increases the range of view angles available for reconstruction, thereby significantly enhancing the quality of Helicon reconstruction. The success of Helicon twist search and reconstruction directly using raw filament images shows another avenue to address the low-twist challenges, although the requirement for long, straight filaments might exclude some helical structures limited to short lengths or being inherently flexible.

Despite the promising results, several limitations of the Helicon method were observed. Firstly, the current implementation of Helicon is computationally intensive, typically taking ∼1 min searching twists in the range of 0.1 to 2° with a 0.1° step and a fixed rise of 4.75 Å for amyloids, thus limiting the search from including additional parameters such as in-plane angles and Y-positions. In the current implementation, the search is parallelized and uses all available CPU cores to speed up the computation. We plan to further speed up the computation with GPU in the future. We also propose that integrating a neural network-based 3D reconstruction approach could offer a solution by optimizing both the helical and angular/positional parameters simultaneously. This will potentially avoid the explicit grid search of the parameters. Additionally, our testing of Helicon on raw filament images was confined to long, straight filaments in raw cryo-EM images. Additional preprocessing steps, including denoising and unbending, due to the inherent noise and distortion in the raw cryo-EM 2D images, will be needed. Addressing these remaining challenges in the future will be critical to extending the applicability of Helicon to more general purposes.

In summary, *helicon denovo3D* utilizes a linear regression model to estimate helical parameters and reconstruct 3D structures from cryo-EM 2D images. The method has proven reliable in estimating helical parameters and producing 3D structures from 2D class average images and raw filament images for various helical structures. Together with the *helicon helicalPitch* program, we show Helicon as a potential solution for helical indexing and *de novo* 3D model for low-twist amyloid structures.

## Resource availability

### Lead contact

Requests for further information and resources should be directed to and will be fulfilled by the lead contact, Wen Jiang.

### Materials availability

This study did not generate new unique reagents.

### Data and code availability

- The test data used in this work are primarily from public databases EMDB and EMPIAR with the accession codes provided in the manuscript.
- The Helicon software reported in this work has been openly developed with its source code and revisions publicly available at https://github.com/jianglab/helicon.

## Acknowledgments

This work was supported in part by NIH grants RF1NS110437, R01AG071177, R21AG081686, Purdue Showalter Scholars award, and Penn State startup fund (WJ). We thank the Jiang lab members for testing the programs and providing feedback during the development of Helicon. We thank Alex Joan Pastor Carattini for proofreading the manuscript.

## Author contributions

Conceptualization, W.J.; Software, W.J., D.L. and X.Z.; Formal Analysis, D.L., X.Z. and W.J.; Writing – Original Draft, D.L.; Writing – Review & Editing, D.L. W.J. and X.Z.; Supervision, W.J.; Funding Acquisition, W.J.

---

**Figure S1:**
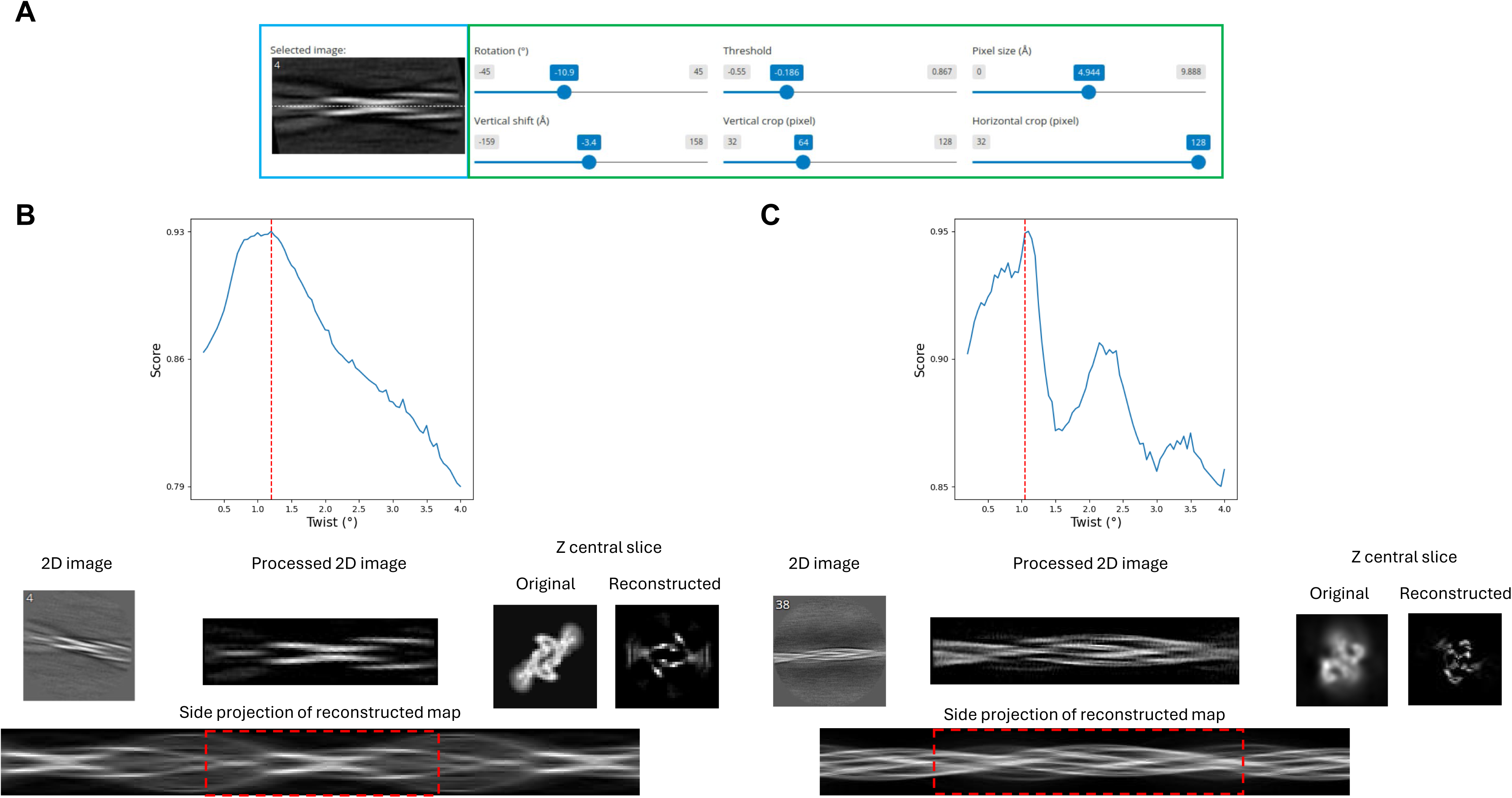
Helicon GUI for in-plane transformation of the input 2D image. The GUI supports both 2D class average images and raw filament images. (A) The GUI to shift and in-plane rotate the 2D image. The green box shows the adjustable parameters. The blue box indicates the transformed image. (B, C) Helicon results using 2D class averages of recombinant tau from EMPIAR-10940 (B) and SF tau from EMPIAR-10230 (C). The in-plane rotation angle, center, and vertical size were adjusted using GUI controls. The red dashed line indicates the ground truth value of the helical twist. The red dashed box represents the reconstructed region corresponding to the original input image.

**Figure S2:**
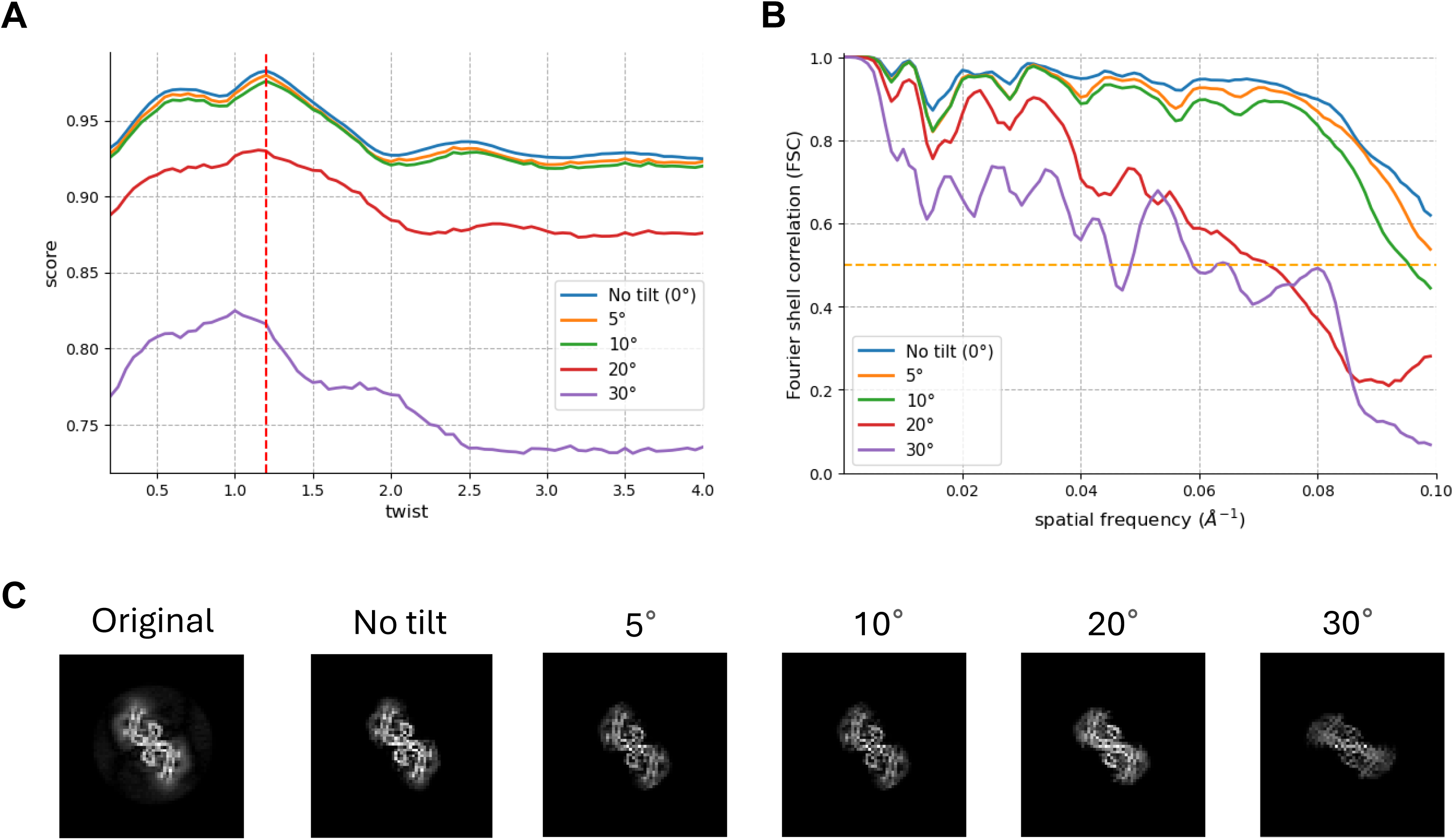
Dependence of Helicon reconstruction quality on out-of-plane tilt angles. Projections with various out-of-plane tilts were generated from the 3D map of PHF tau (EMD-23894). (A) Helicon score curves. (B) FSC of the ground truth density and the Helicon 3D reconstructions. (C) The central Z-slice of the Helicon reconstructions.

**Figure S3:**
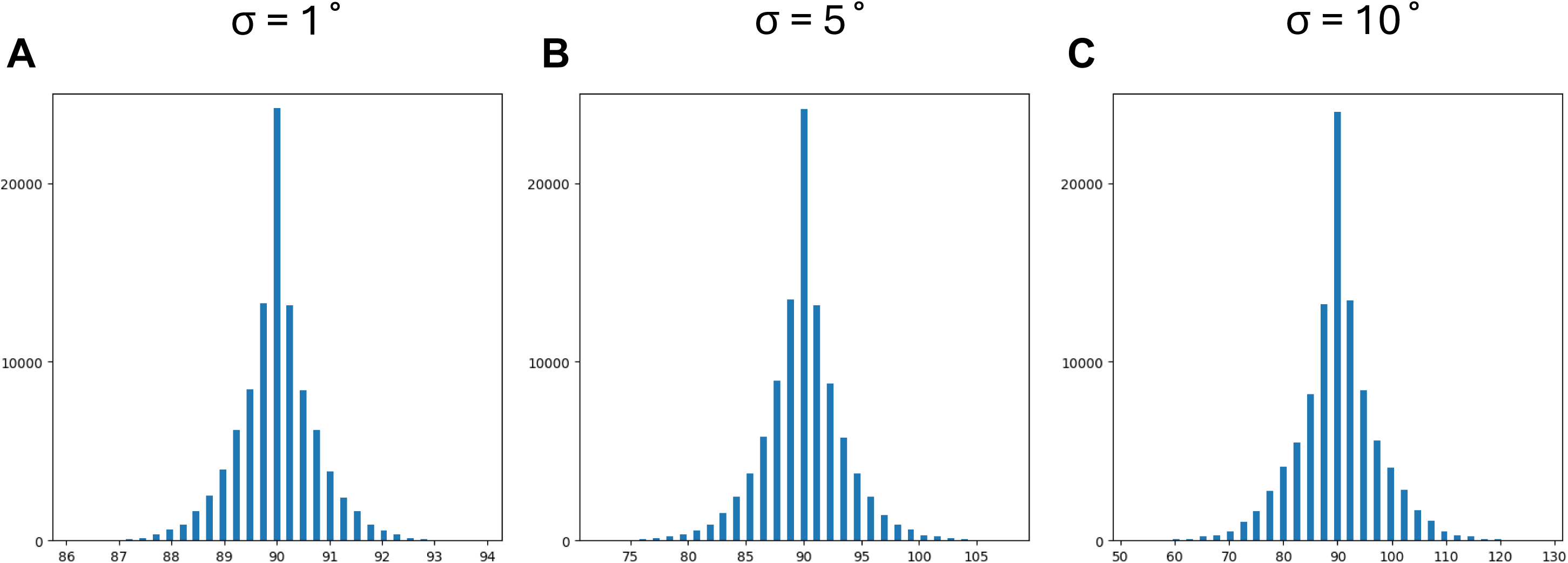
Simulation of the distribution of filament tilt angles under various levels of grid tilt angles. Histogram plots showing the distribution of filament tilt angles for different standard deviations (σ) of grid tilt angles: 1° (A), 5° (B), and 10° (C) assuming random in-plane angles of filaments 10000 filaments in 1000 micrographs were used in the simulations.

**Figure S4:**
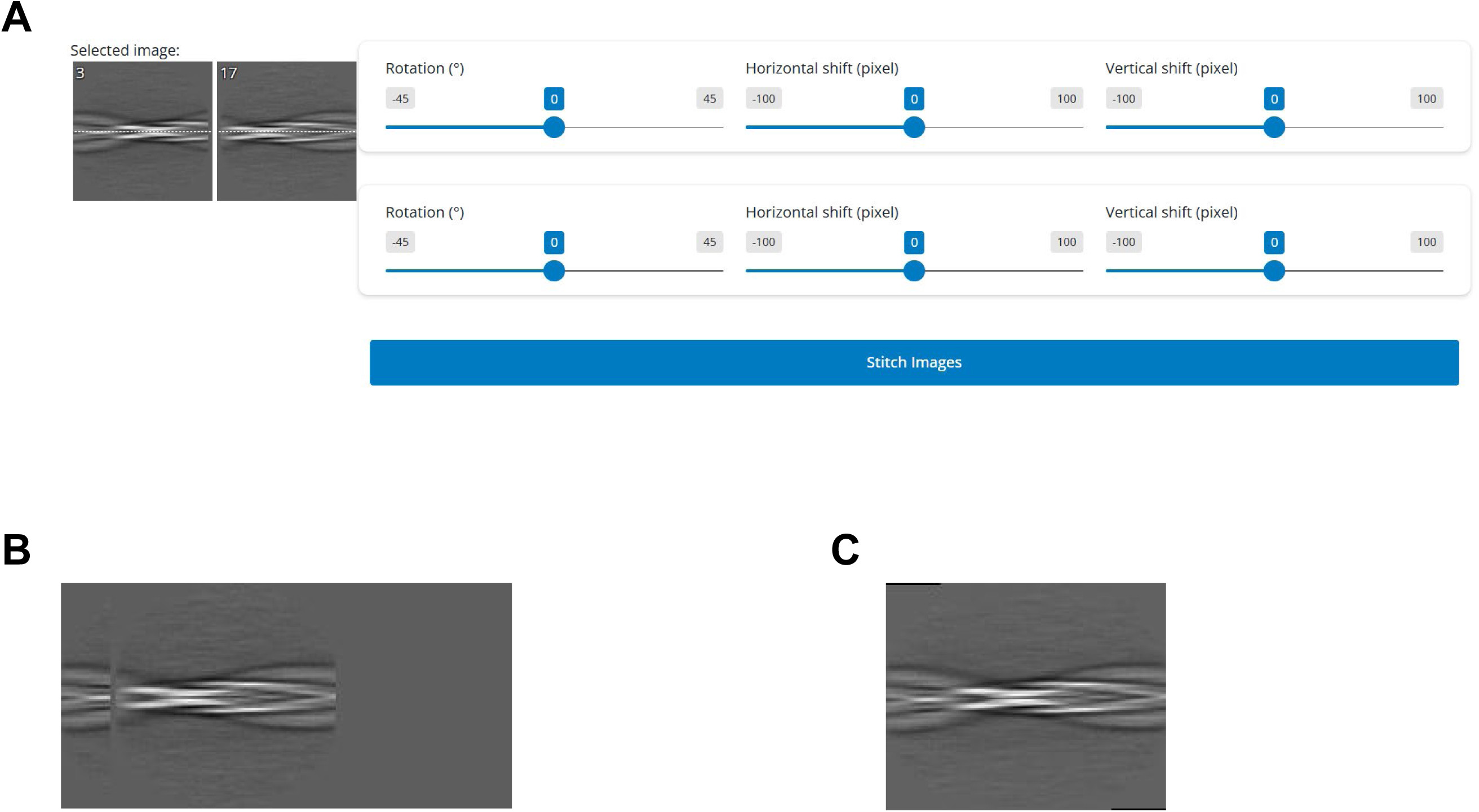
Helicon GUI for image stitching. (A) The adjustable parameters for image stitching. (B) An example of two superimposed images transformed based on the stitching parameters. (C) An example of a stitched image.

**Figure S5:**
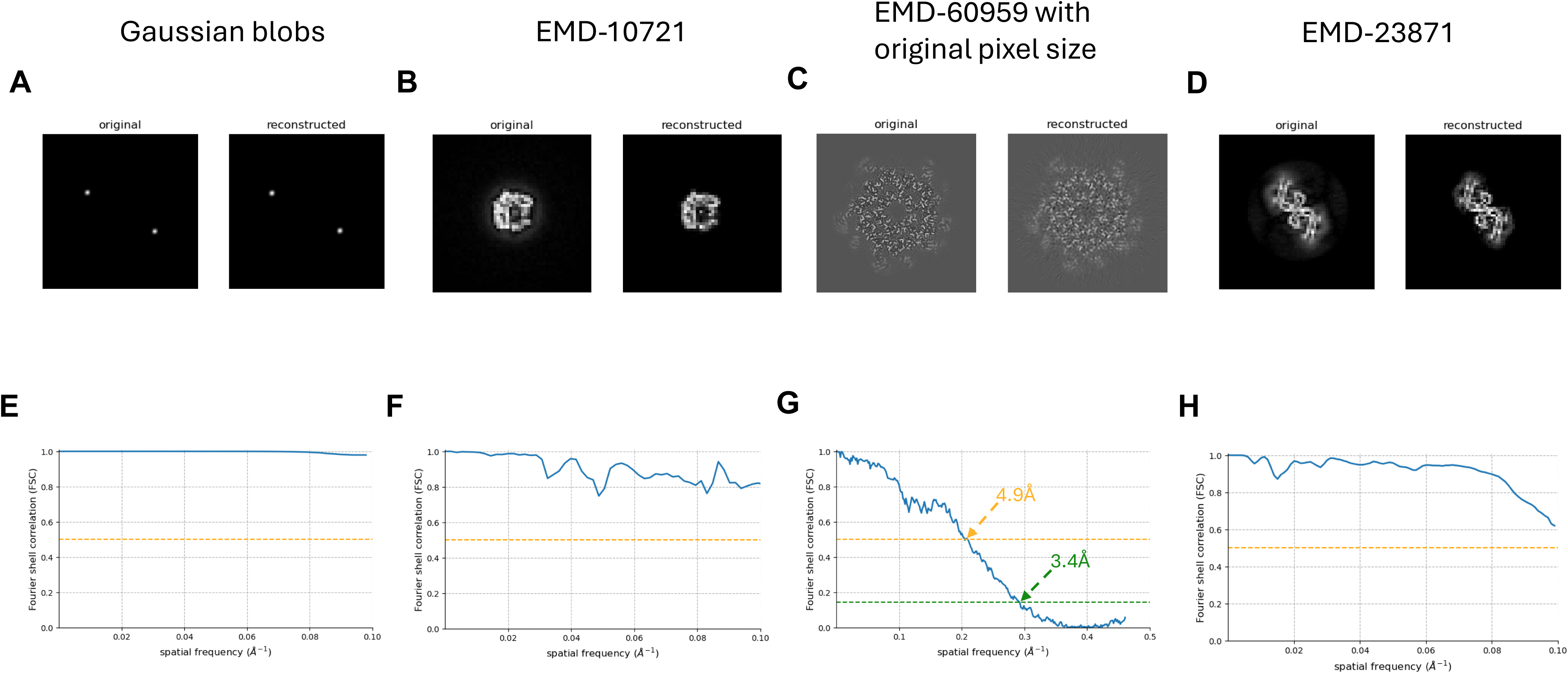
Comparison of Helicon 3D reconstructions and the ground truth density maps. (A-D) The central slice of the ground truth density maps (left) and the corresponding Helicon 3D reconstructions (right). (E-H) The corresponding FSC curves between the ground truth density maps and the Helicon 3D reconstructions. The FSC thresholds of 0.143 and 0.5 are shown as green and yellow lines, respectively.

**Figure S6:**
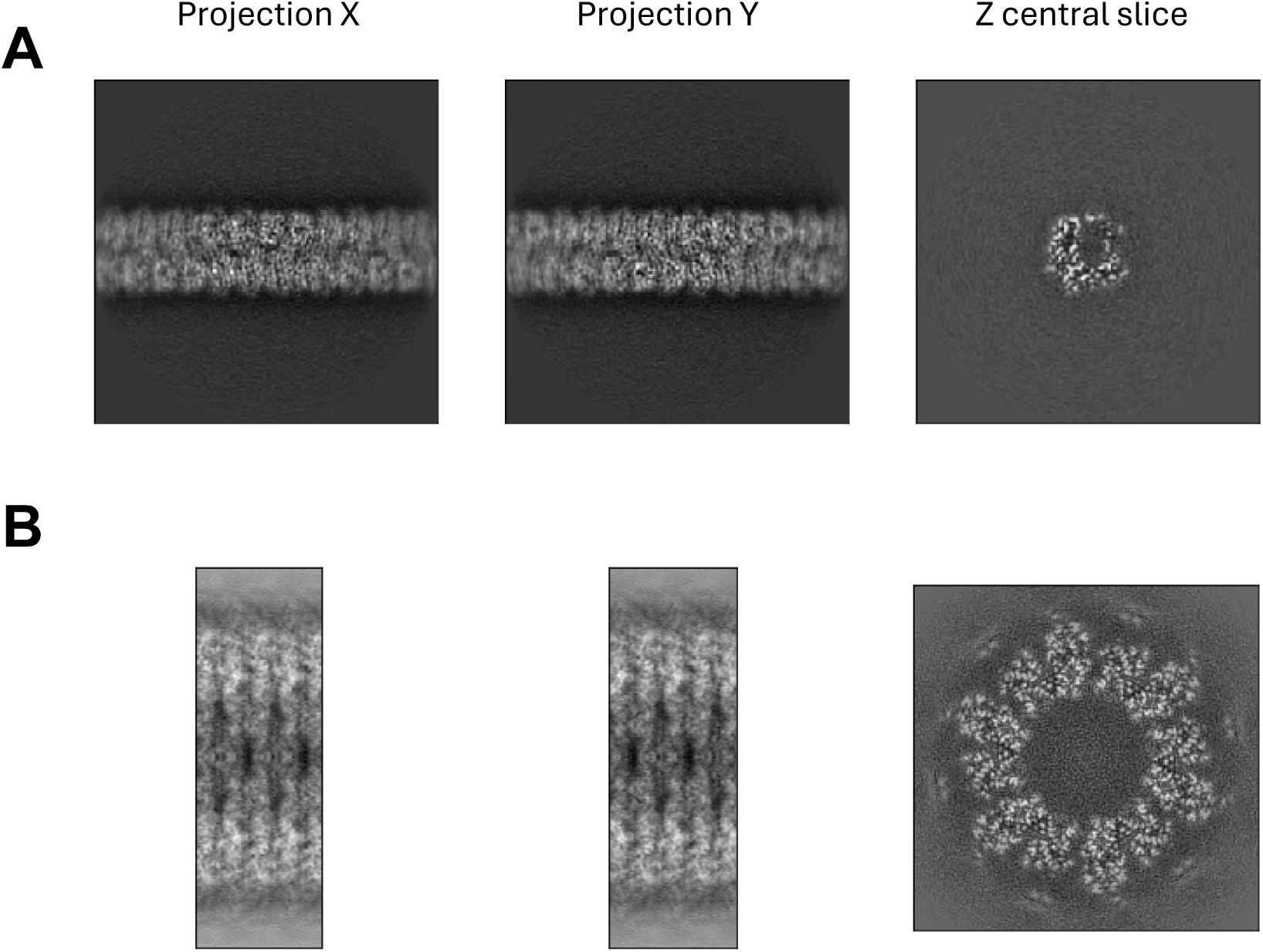
The ground truth density of the two non-amyloid helical structures. Shown are the side projections and central z slice of high-resolution 3D structures solved from cryo-EM datasets of bacteria pili (A) and VipA/B from EMPIAR-10019 (B). The class averages of these two datasets were used in the Helicon tests shown in Figure 3.

**Figure S7:**
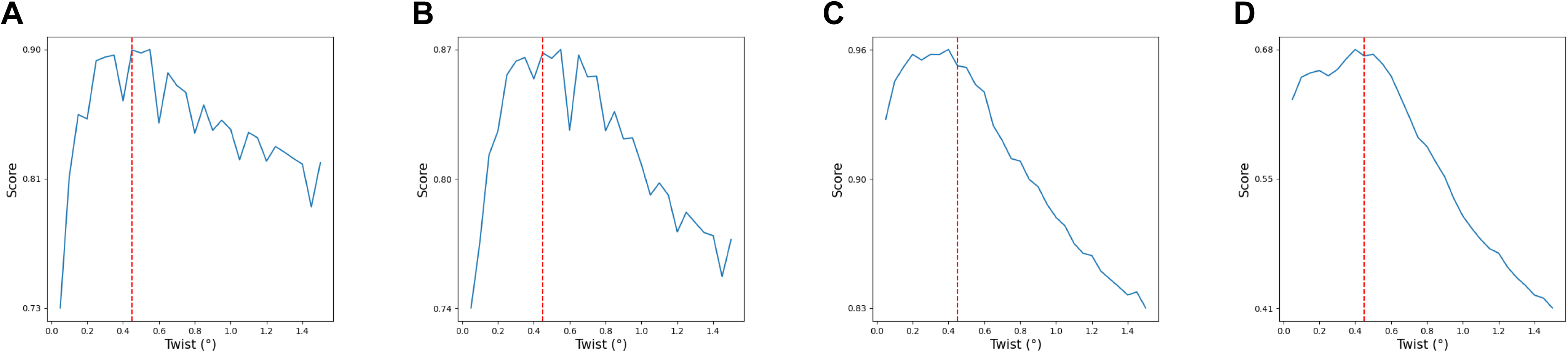
Helicon score curves with different regularization weights. The sample is type 3b amyloid β with a low twist of 0.46°. Different ElasticNet regularization weights, 1e-7 (A), 1e-5 (B), 1e-3 (C), and 1 (D) were shown. The red dashed line indicates the ground truth value of the helical twist.

**Figure S8:**
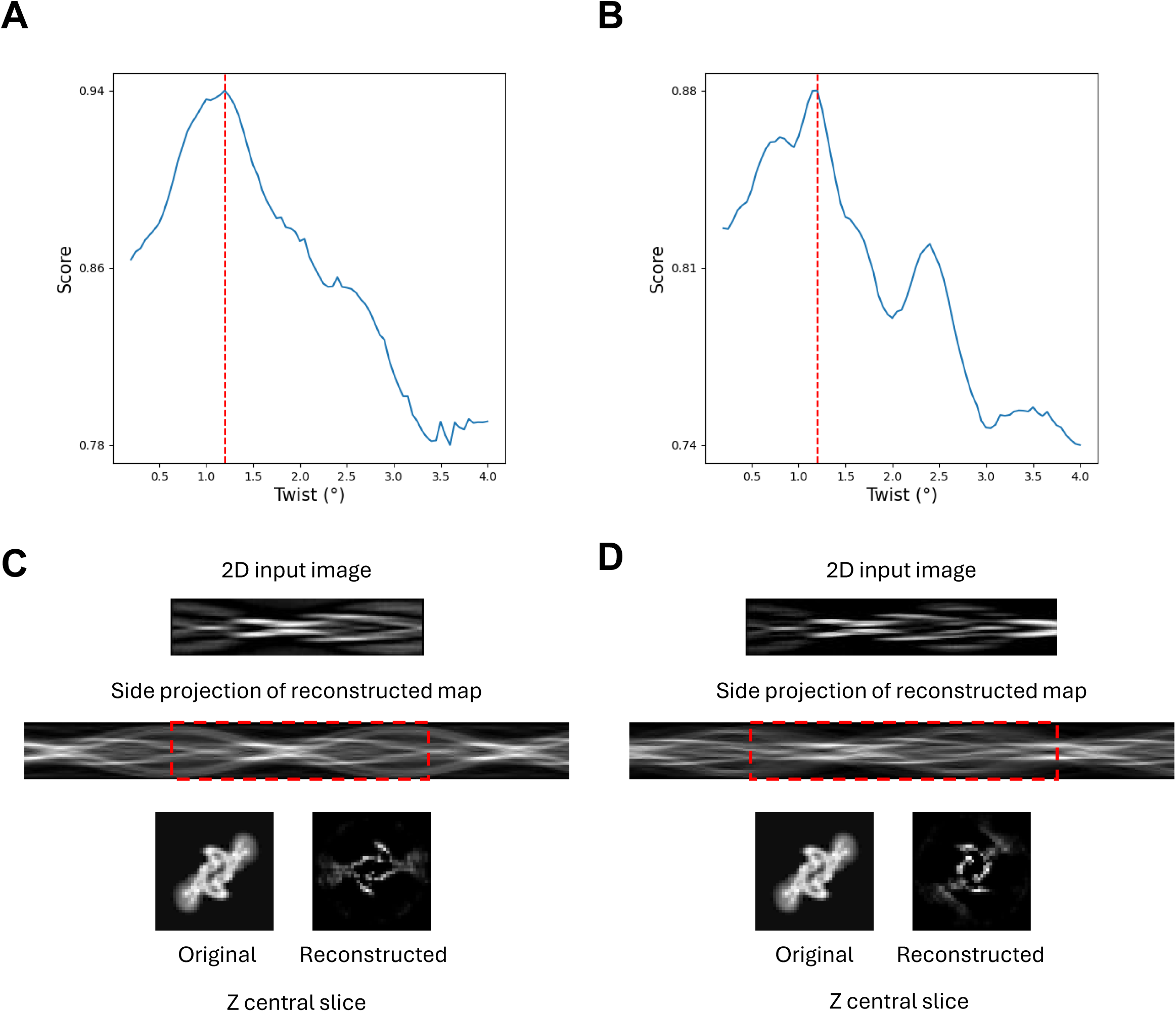
Example of Helicon reconstruction of stitched images. The example shown here uses 2D class average images in the EMPIAR-10940 dataset. (A, B) Helicon score curves for images with a stitched segment length of 800 Å (A) and 1000 Å (B). (C, D) Side projects and Z central slice of Helicon reconstructions using the optimal twist values shown in A and B, respectively. The red dashed lines indicate the ground truth value of the helical twist. The red dashed boxes highlight the regions of Helicon reconstruction corresponding to the original input images.

**Figure S9:**
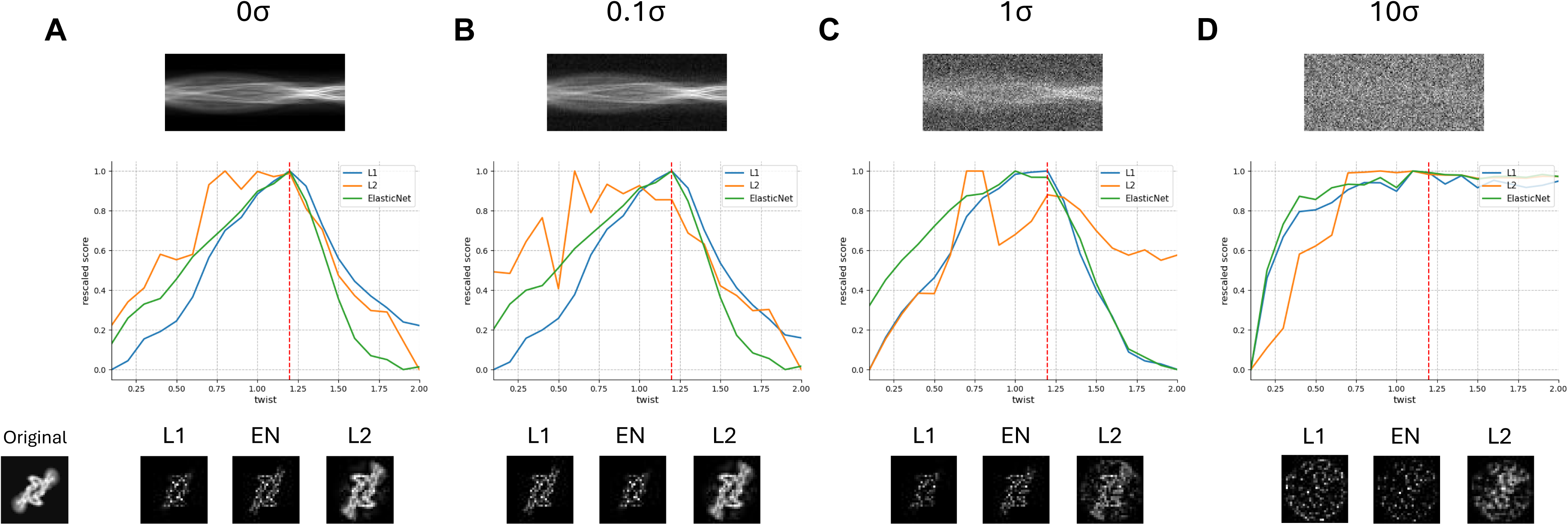
Helicon score curves and reconstruction quality for images with varying SNR and types of regularizations. The sample consists of projections of EMD-14046 (recombinant tau) with various levels of added noise. The figures illustrate the score curves obtained using L1, L2, and ElasticNet (i.e. equal weights of L1 and L2) regularization, along with the corresponding reconstructions using the ground truth twist of −1.2°. The noise level is represented by the multiples of the variance of the signal (σ) at 0 σ (A), 0.1 σ (B), 1 σ (C), and 10 σ (D), respectively.

**Figure S10:**
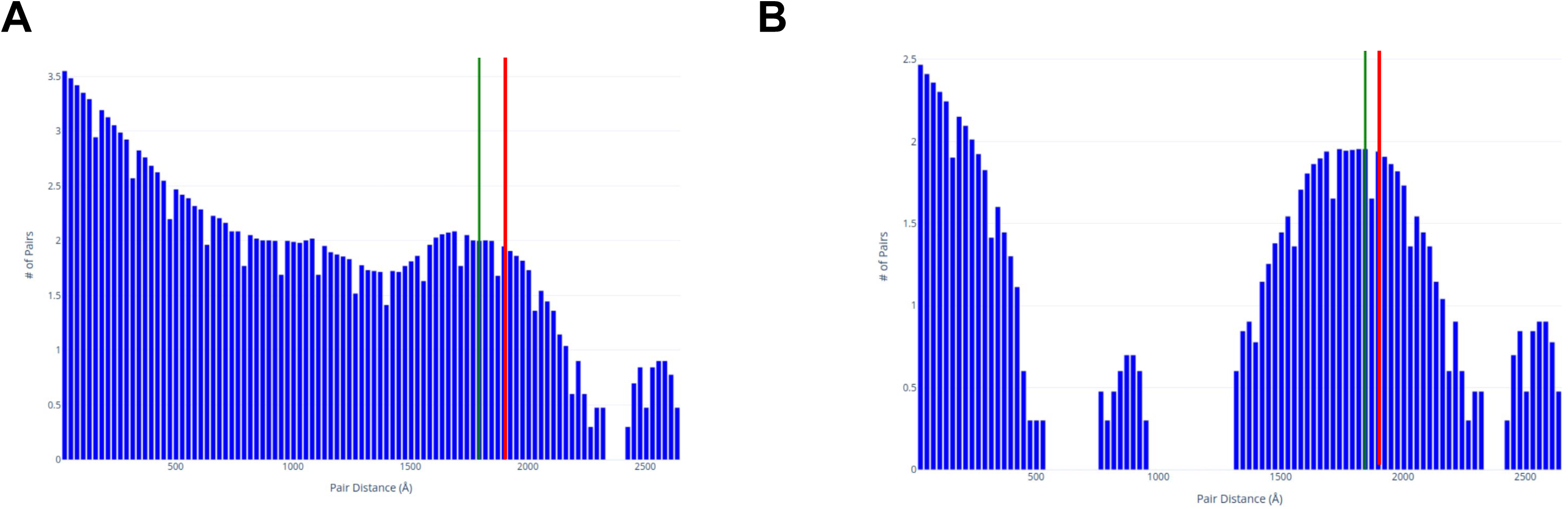
Comparison of *HelicalPitch* histogram plots using all filaments and long filaments only. The data used in this example is the type 3b amyloid β filaments in the Down syndrome patient samples having a low twist of 0.46° and a pitch of 3,725 Å. (A) Using all filaments. (B) Using only filaments longer than 2500 Å.

**Figure S11:**
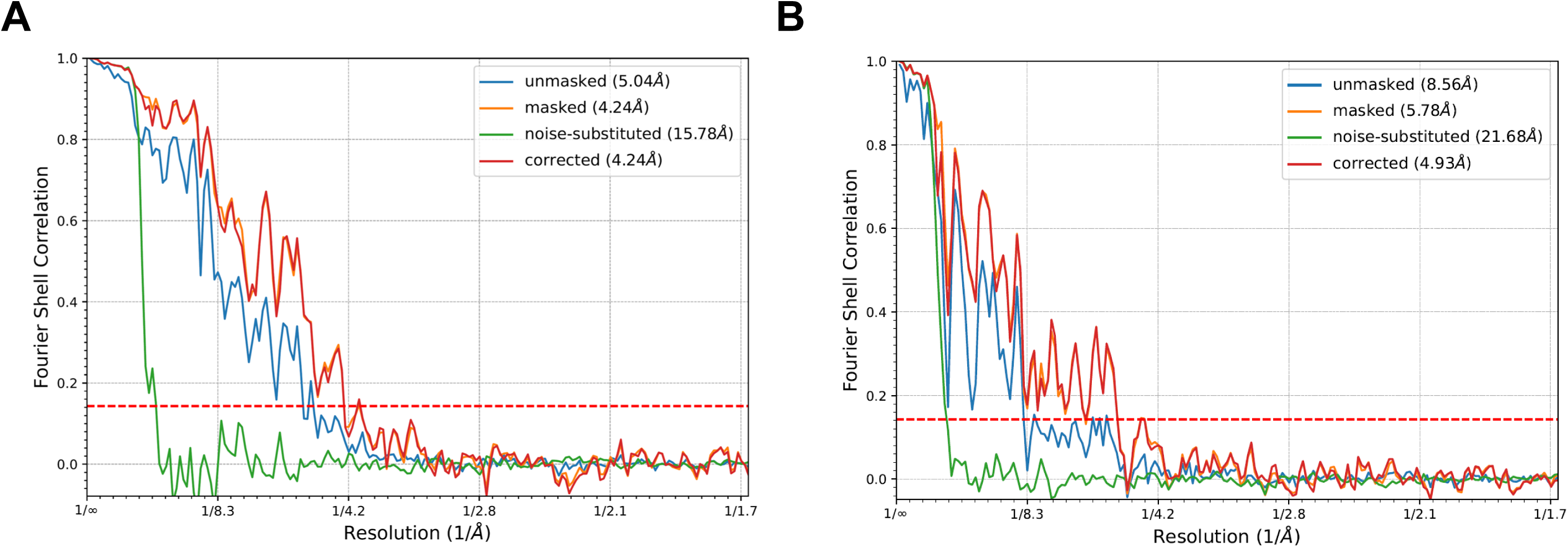
The FSC curves of the refinement results for the 2^nd^ type of tau filament in EMPIAR-10940. (A) from all 44 filaments. (B) from only the two longest filaments.

## References

1. Watson, J.D., and Crick, F.H.C. (1953). Molecular Structure of Nucleic Acids: A Structure for Deoxyribose Nucleic Acid. Nature 171, 737–738. 10.1038/171737a0.

2. Chakraborty, S., Jasnin, M., and Baumeister, W. (2020). Three-dimensional organization of the cytoskeleton: A cryo-electron tomography perspective. Protein Sci. Publ. Protein Soc. 29, 1302–1320. 10.1002/pro.3858.

3. Fäßler, F., Javoor, M.G., and Schur, F.K. (2023). Deciphering the molecular mechanisms of actin cytoskeleton regulation in cell migration using cryo-EM. Biochem. Soc. Trans. 51, 87–99. 10.1042/BST20220221.

4. Conners, R., León-Quezada, R.I., McLaren, M., Bennett, N.J., Daum, B., Rakonjac, J., and Gold, V.A.M. (2023). Cryo-electron microscopy of the f1 filamentous phage reveals insights into viral infection and assembly. Nat. Commun. 14, 2724. 10.1038/s41467-023-37915-w.

5. González, B., Li, D., Li, K., Wright, E.T., Hardies, S.C., Thomas, J.A., Serwer, P., and Jiang, W. (2021). Structural Studies of the Phage G Tail Demonstrate an Atypical Tail Contraction. Viruses 13, 2094. 10.3390/v13102094.

6. Fernandez, A., Hoq, M.R., Hallinan, G.I., Li, D., Bharath, S.R., Vago, F.S., Zhang, X., Ozcan, K.A., Newell, K.L., Garringer, H.J., et al. (2024). Cryo-EM structures of amyloid-β and tau filaments in Down syndrome. Nat. Struct. Mol. Biol. 31, 903–909. 10.1038/s41594-024-01252-3.

7. Fitzpatrick, A.W.P., Falcon, B., He, S., Murzin, A.G., Murshudov, G., Garringer, H.J., Crowther, R.A., Ghetti, B., Goedert, M., and Scheres, S.H.W. (2017). Cryo-EM structures of tau filaments from Alzheimer’s disease. Nature 547, 185–190. 10.1038/nature23002.

8. Lövestam, S., Li, D., Wagstaff, J.L., Kotecha, A., Kimanius, D., McLaughlin, S.H., Murzin, A.G., Freund, S.M.V., Goedert, M., and Scheres, S.H.W. (2024). Disease-specific tau filaments assemble via polymorphic intermediates. Nature 625, 119–125. 10.1038/s41586-023-06788-w.

9. Zhu, K.-F., Yuan, C., Du, Y.-M., Sun, K.-L., Zhang, X.-K., Vogel, H., Jia, X.-D., Gao, Y.-Z., Zhang, Q.-F., Wang, D.-P., et al. (2023). Applications and prospects of cryo-EM in drug discovery. Mil. Med. Res. 10, 10. 10.1186/s40779-023-00446-y.

10. Yip, K.M., Fischer, N., Paknia, E., Chari, A., and Stark, H. (2020). Atomic-resolution protein structure determination by cryo-EM. Nature 587, 157–161. 10.1038/s41586-020-2833-4.

11. De Rosier, D.J., and Klug, A. (1968). Reconstruction of Three Dimensional Structures from Electron Micrographs. Nature 217, 130–134. 10.1038/217130a0.

12. Wang, F., Gnewou, O., Solemanifar, A., Conticello, V.P., and Egelman, E.H. (2022). Cryo-EM of Helical Polymers. Chem. Rev., 10.1021/acs.chemrev.1c00753. 10.1021/acs.chemrev.1c00753.

13. Diaz, R., Rice, W., J., and Stokes, D.L. (2010). Chapter Five - Fourier–Bessel Reconstruction of Helical Assemblies. In Methods in Enzymology Cryo-EM, Part B: 3-D Reconstruction., G. J. Jensen, ed. (Academic Press), pp. 131–165. 10.1016/S0076-6879(10)82005-1.

14. Egelman, E.H. (2010). Reconstruction of Helical Filaments and Tubes. Methods Enzymol. 482, 167–183. 10.1016/S0076-6879(10)82006-3.

15. He, S., and Scheres, S.H.W. (2017). Helical reconstruction in RELION. J. Struct. Biol. 198, 163–176. 10.1016/j.jsb.2017.02.003.

16. Egelman, E.H. (2014). Ambiguities in helical reconstruction. eLife 3, e04969. 10.7554/eLife.04969.

17. Scheres, S.H.W. (2020). Amyloid structure determination in RELION-3.1. Acta Crystallogr. Sect. Struct. Biol. 76, 94–101. 10.1107/S2059798319016577.

18. Yan, R., Venkatakrishnan, S.V., Liu, J., Bouman, C.A., and Jiang, W. (2019). MBIR: A cryo-ET 3D reconstruction method that effectively minimizes missing wedge artifacts and restores missing information. J. Struct. Biol. 206, 183–192.

19. Zou, H., and Hastie, T. (2005). Regularization and Variable Selection Via the Elastic Net. J. R. Stat. Soc. Ser. B Stat. Methodol. 67, 301–320. 10.1111/j.1467-9868.2005.00503.x.

20. Li, D., and Jiang, W. (2023). Classification of helical polymers with deep-learning language models. J. Struct. Biol. 215, 108041. 10.1016/j.jsb.2023.108041.

21. Zyla, D.S., Wiegand, T., Bachmann, P., Zdanowicz, R., Giese, C., Meier, B.H., Waksman, G., Hospenthal, M.K., and Glockshuber, R. (2024). The assembly platform FimD is required to obtain the most stable quaternary structure of type 1 pili. Nat. Commun. 15, 3032. 10.1038/s41467-024-47212-9.

22. Hallinan, G.I., Hoq, M.R., Ghosh, M., Vago, F.S., Fernandez, A., Garringer, H.J., Vidal, R., Jiang, W., and Ghetti, B. (2021). Structure of Tau filaments in Prion protein amyloidoses. Acta Neuropathol. (Berl.) 142, 227–241. 10.1007/s00401-021-02336-w.

23. Lövestam, S., and Scheres, S.H.W. (2022). High-throughput cryo-EM structure determination of amyloids. Faraday Discuss. 240, 243–260. 10.1039/D2FD00034B.

24. Kudryashev, M., Wang, R.Y.-R., Brackmann, M., Scherer, S., Maier, T., Baker, D., DiMaio, F., Stahlberg, H., Egelman, E.H., and Basler, M. (2015). Structure of the Type VI Secretion System Contractile Sheath. Cell 160, 952–962. 10.1016/j.cell.2015.01.037.

25. Lövestam, S., Schweighauser, M., Matsubara, T., Murayama, S., Tomita, T., Ando, T., Hasegawa, K., Yoshida, M., Tarutani, A., Hasegawa, M., et al. (2021). Seeded assembly in vitro does not replicate the structures of α-synuclein filaments from multiple system atrophy. FEBS Open Bio 11, 999–1013. 10.1002/2211-5463.13110.

26. Yang, Y., Arseni, D., Zhang, W., Huang, M., Lövestam, S., Schweighauser, M., Kotecha, A., Murzin, A.G., Peak-Chew, S.Y., Macdonald, J., et al. (2022). Cryo-EM structures of amyloid-β 42 filaments from human brains. Science. 10.1126/science.abm7285.

27. Falcon, B., Zhang, W., Schweighauser, M., Murzin, A.G., Vidal, R., Garringer, H.J., Ghetti, B., Scheres, S.H.W., and Goedert, M. (2018). Tau filaments from multiple cases of sporadic and inherited Alzheimer’s disease adopt a common fold. Acta Neuropathol. (Berl.) 136, 699–708. 10.1007/s00401-018-1914-z.

28. Zhang, W., Falcon, B., Murzin, A.G., Fan, J., Crowther, R.A., Goedert, M., and Scheres, S.H. (2019). Heparin-induced tau filaments are polymorphic and differ from those in Alzheimer’s and Pick’s diseases. eLife 8, e43584. 10.7554/eLife.43584.

29. Lövestam, S., Koh, F.A., van Knippenberg, B., Kotecha, A., Murzin, A.G., Goedert, M., and Scheres, S.H. (2022). Assembly of recombinant tau into filaments identical to those of Alzheimer’s disease and chronic traumatic encephalopathy. eLife 11, e76494. 10.7554/eLife.76494.

30. Zheng, S.Q., Palovcak, E., Armache, J.-P., Verba, K.A., Cheng, Y., and Agard, D.A. (2017). MotionCor2: anisotropic correction of beam-induced motion for improved cryo-electron microscopy. Nat. Methods 14, 331–332. 10.1038/nmeth.4193.

31. Rohou, A., and Grigorieff, N. (2015). CTFFIND4: Fast and accurate defocus estimation from electron micrographs. J. Struct. Biol. 192, 216–221. 10.1016/j.jsb.2015.08.008.

32. Huber, S.T., Kuhm, T., and Sachse, C. (2018). Automated tracing of helical assemblies from electron cryo-micrographs. J. Struct. Biol. 202, 1–12. 10.1016/j.jsb.2017.11.013.

33. Tang, G., Peng, L., Baldwin, P.R., Mann, D.S., Jiang, W., Rees, I., and Ludtke, S.J. (2007). EMAN2: An extensible image processing suite for electron microscopy. J. Struct. Biol. 157, 38–46. 10.1016/j.jsb.2006.05.009.

34. Zukić, D., Jackson, M., Dimiduk, D., Donegan, S., Groeber, M., and McCormick, M. (2021). ITKMontage: A Software Module for Image Stitching. Integrating Mater. Manuf. Innov. 10, 115–124. 10.1007/s40192-021-00202-x.

35. Kimanius, D., Jamali, K., Wilkinson, M.E., Lövestam, S., Velazhahan, V., Nakane, T., and Scheres, S.H.W. (2024). Data-driven regularization lowers the size barrier of cryo-EM structure determination. Nat. Methods 21, 1216–1221. 10.1038/s41592-024-02304-8.

36. Tibshirani, R. (1996). Regression Shrinkage and Selection Via the Lasso. J. R. Stat. Soc. Ser. B Methodol. 58, 267–288. 10.1111/j.2517-6161.1996.tb02080.x.

37. Hastie, T., Tibshirani, R., and Friedman, J. (2009). Linear Methods for Regression. In The Elements of Statistical Learning: Data Mining, Inference, and Prediction, T. Hastie, R. Tibshirani, and J. Friedman, eds. (Springer), pp. 43–99. 10.1007/978-0-387-84858-7_3.

38. Tan, Y.Z., Baldwin, P.R., Davis, J.H., Williamson, J.R., Potter, C.S., Carragher, B., and Lyumkis, D. (2017). Addressing preferred specimen orientation in single-particle cryo-EM through tilting. Nat. Methods 14, 793–796. 10.1038/nmeth.4347.

